# Genome-scale conserved molecular principles of mRNA half-life regulation

**DOI:** 10.1101/2020.02.17.952267

**Authors:** Sudipto Basu, Saurav Mallik, Suman Hait, Sudip Kundu

## Abstract

Precise control of protein and mRNA degradation is essential for cellular metabolism and homeostasis. Controlled and specific degradation of both molecular species necessitates their engagements with the respective degradation machineries; this engagement involves a disordered/unstructured segment of the substrate traversing the degradation tunnel of the machinery and accessing the catalytic sites. Here, we report that mRNAs comprising longer terminal and/or internal unstructured segments have significantly shorter half-lives; the lengths of the 5′ terminal, 3′ terminal and internal unstructured segments that affect mRNA half-life are compatible with molecular structures of the 5′ exo- 3′ exo- and endo-ribonuclease machineries. Sequestration into ribonucleoprotein complexes elongates mRNA half-life, presumably by burying ribonuclease engagement sites under oligomeric interfaces. After gene duplication, differences in terminal unstructured lengths, proportions of internal unstructured segments and oligomerization modes result in significantly altered half-lives of paralogous mRNAs. Side-by-side comparison of molecular principles underlying controlled protein and mRNA degradation unravels their remarkable mechanistic similarities, and suggests how the intrinsic structural features of the two molecular species regulate their half-lives on genome-scale and during evolution.

## Introduction

Quality control of all the constituent molecular machines is the key to sustain the living engine we call a cell. A plethora of surveillance pathways have evolved at different kingdoms of life to promote and to sustain the accurate designing of all the cellular macromolecules. For DNA, which is replicated only once during a cell cycle, elegant repair mechanisms have evolved to orchestrate the quality control of genetic information with chromosome segregation and cell cycle progression (Hustedt & Durocher, 2016). For RNA and proteins, which are regularly synthesized, a myriad of enzyme machineries has evolved to prompt the degradation of the aberrant species and to execute their controlled hydrolysis as they get damaged during their lifetime (Houseley & Tollervey, 2009; Bhattacharyya *et al*, 2014). A precise balance between the synthesis of nascent copies and degradation of the damaged ones maintains the cellular homeostasis and assigns each protein and RNA molecule a characteristic half-life (Belle *et al*, 2006; Mathieson *et al*, 2018; Price *et al*, 2010; Wang *et al*, 2002; Narsai *et al*, 2007; Raghavan *et al*, 2002; Eser *et al*, 2016)

Despite the immense compositional, structural and functional diversity of protein and RNA molecules within the cell and among organisms, their controlled degradations, in all kingdoms of life, exhibit remarkably similar mechanistic principles (Makino *et al*, 2013b; Houseley & Tollervey, 2009; Cromm & Crews, 2017). In eukaryotic cells, degradation of both molecular species involves surveillance and proofreading pathways recognizing the damaged copies and festooning them with a degradation signal (poly(A/U) tag (Slomovic *et al*, 2010; West *et al*, 2006; Bresson *et al*, 2015; Mullen & Marzluff, 2008) and 5′ decapping (Mullen & Marzluff, 2008; Franks & Lykke-Andersen, 2008; Coller & Parker, 2004) for mRNAs, and poly-ubiquitin tag for proteins (Finley, 2009; Thrower *et al*, 2000). Protease and RNase machineries then recognize these aberrant substrates based on these tags, mechanically unfold them with the help of ATP-dependent and/or ATP-independent cofactors and finally hydrolyze them into small oligo-peptide/-nucleotide fragments (Bhattacharyya *et al*, 2014; Makino *et al*, 2013b; Houseley & Tollervey, 2009).

Both protein and RNA degradation in the cell follow first order kinetics, depending on a rate-determining initial step: substrate engagement with the protease/RNase machinery (Goldberg & Dice, 1974; Schimke & Doyle, 1970; Laalami *et al*, 2014). Protein degradation in eukaryotic cells is predominantly mediated by a barrel-shaped self-assembling machine called proteasome that can engage with any disordered region of the substrate of sufficient length (∼30 amino acid for terminal, ∼40 amino acid for internal) and initiate degradation (Ciechanover, 2005; Bhattacharyya *et al*, 2014; van der Lee *et al*, 2014). Prevalence of one single, highly conserved machinery confers the advantage that understanding how it works leads to a systematic understanding of the overall process in a multitude of organisms. Consequently, molecular factors influencing protein degradation have been extensively explored on a genome scale, and in multiple organisms (van der Lee *et al*, 2014; Fishbain *et al*, 2015; Mallik & Kundu, 2018). RNA degradation, on the other hand, involves three major classes of RNases: endonucleases that cut RNA internally, 5′ exoribonucleases that hydrolyze RNA from the 5′ end, and 3′ exoribonucleases that degrade RNA from the 3′ end (Houseley & Tollervey, 2009; Hui *et al*, 2014). Genomes of most organisms encode multiple enzymes of each class, often with overlapping activities and a plethora of common target substrates, which makes the overall process very difficult to understand systematically (Houseley & Tollervey, 2009; Hui *et al*, 2014). As a result, to this date exploration of molecular factors influencing RNA (and especially messenger RNA, mRNA) degradation on a genome scale remains limited to finding the degradation signals (Slomovic *et al*, 2010; West *et al*, 2006; Bresson *et al*, 2015; Mullen & Marzluff, 2008; Franks & Lykke-Andersen, 2008; Coller & Parker, 2004) and degradation-promoting or hindering sequence motifs (Tomecki & Dziembowski, 2010; Cheng *et al*, 2017; Yang *et al*, 2003; Geisberg *et al*, 2014; Geissler & Grimson, 2016). But a systematic understanding of how the structural attributes of the RNase machineries and their substrates influence mRNA half-life on a genome scale and during evolution remains elusive.

Here, we exploited (i) the experimental half-life data of *Saccharomyces cerevisiae* mRNAs, (Data S1) (ii) their experimentally derived secondary structures as well as protein-binding data, (iii) extensive characterization of their coding (CDS) and 5′ and 3′ untranslated regions (UTRs), (Data S1) along with (iv) high-resolution X-ray crystallographic structures of the RNase machines (Table S1) – to develop a comprehensive theory demonstrating how the intrinsic structural attributes of the RNase machines and their substrates influence mRNA half-life on a genome scale. Despite the enormous variation in sequence, structure and function of mRNAs on a genome scale, and that of their degradation machineries, simple molecular principles tune transcript half-lives across the genome and during evolution, similar to that of proteins.

## Results

To investigate the relationship between the structural attributes of mRNA transcripts and their *in vivo* half-lives, we began with comparing the known mechanistic principles of protein and mRNA degradation, to gain insight about their similarities. The major protease machinery, proteasome, is a barrel-shaped molecular machine, with its catalytic sites accessible through a ∼70 Å long narrow internal tunnel. Protein substrates that exhibit ∼30 residues long intrinsically disordered regions at their termini or ∼40 residues long intrinsically disordered regions in the middle (Lobanov *et al*, 2010; Uversky, 2013; van der Lee *et al*, 2014) can potentially traverse this tunnel and access the catalytic sites. This geometrical constraint of enzyme-substrate engagement tunes protein half-life in a way that proteins comprising >30 residues terminal (**Fig. 1A**) and/or >40 residues internal disordered regions (**Fig. 1B**) exhibit significantly shorter half-life than proteins without these features (van der Lee *et al*, 2014). A careful survey of the existing literature hints that similar mechanistic principles influence mRNA half-life as well. On one hand, crystal structure data showed that mRNA degradation machineries exhibit ‘molecular cage’-like shapes, with their catalytic sites accessible through narrow internal tunnels (**Fig. 1 C–G**) (except for endonuclease degradation where the catalytic sites are positioned on the surface (Bonneau *et al*, 2009)). On the other hand, experimental genome-wide mRNA secondary structure measurements (Rouskin *et al*, 2014; Kertesz *et al*, 2010) as well as computational predictions (Shabalina *et al*, 2006) depicted that mRNAs tend to be more unstructured (presence of higher proportion of single stranded region) at their termini than in the middle (**Fig 2A**). To systematically understand how the geometrical constraints of enzyme-substrate engagement influence mRNA half-life on a genome-scale and in evolution, we identified the degradation tunnels in different RNase machines and estimated the lengths of terminal unstructured regions for each mRNA substrates. Combining these with experimental mRNA half-life data, we performed a rigorous statistical comparison.

**Figure 1.**
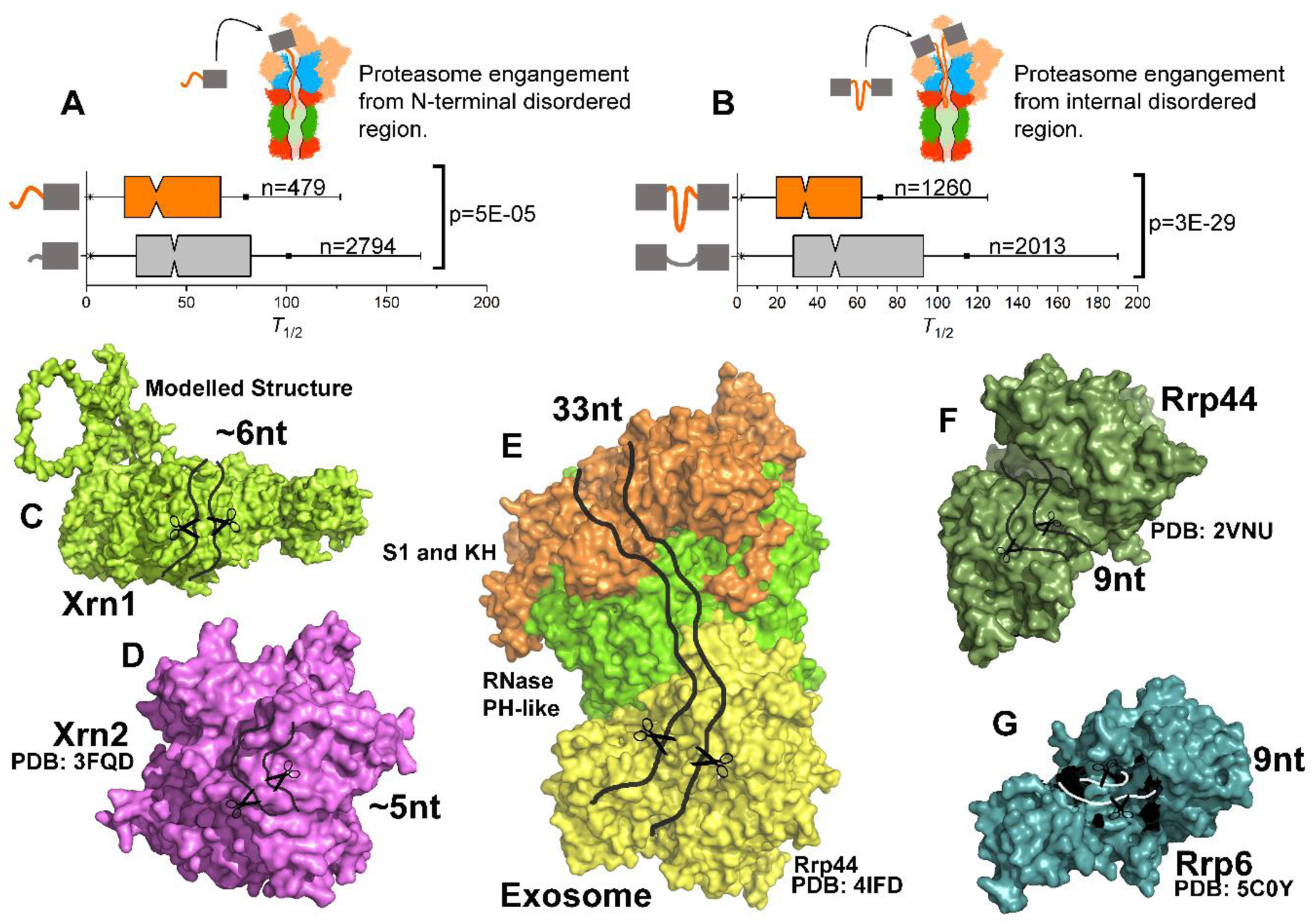
RNase machineries exhibit catalytic sites accessible through narrow internal tunnels. **(A–B)** Schematic diagrams of proteins engaging with proteasome through their **(A)** terminal and **(B)** internal disordered regions. **Bottom.** Proteins were classified based on the lengths of their **(A)** terminal and **(B)** internal disordered segments: long (TDR > 30 aa, and IDR > 40 aa, orange) and short (TDR ≤ 30 aa, and IDR ≤ 40 aa, grey). Mann-Whitney U tests were performed to test whether the distributions significantly differ, *p*-values are mentioned. These two panels are generated following ref. (van der Lee *et al*, 2014). (**C–G**) The molecular diagrams of different mRNA machineries (surface representation), with their degradation tunnels highlighted schematically. The catalytic site locations are schematically marked as scissors. The machineries depicted as following: progressive 5′→3′ exoribonuclease Xrn1 **(C)** and Xrn2 **(D)**, the Exo-10 (comprising RNase PH-like subunits, green, and S1 and KH ring, orange) associated with progressive 3′→5′ exoribonuclease Rrp44 (yellow) **(E)**, Rrp44 acting alone **(F)**, distributive exoribonuclease Rrp6 acting alone **(G)**.

**Figure 2.**
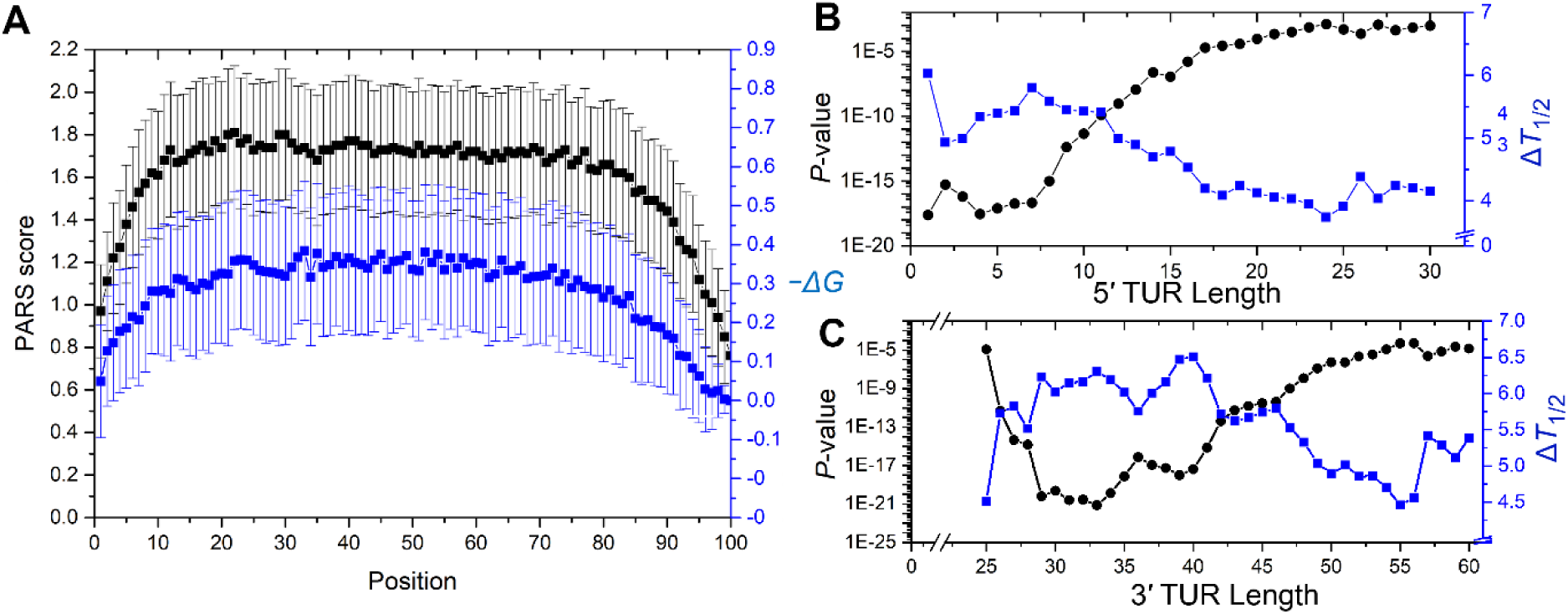
The effects of terminal unstructured region lengths on *S. cerevisiae* mRNA half-life. **(A)** PARS scores (black) and RNAfold-predicted −ΔG values (blue), averaged over all the transcripts (n=3000) used in our analysis, are plotted, where each transcript length is scaled within the range 1-100. For both PARS and −ΔG, the respective standard deviation values (shown as error bars) are scaled down five-fold, for the clarity of presentation. **(B–C)** For different **(B)** 5′ TUR and **(C)** 3′ TUR length thresholds, yeast transcripts (n=2854) were classified into two groups: short (those that exhibit TUR lengths ≤ threshold) and long (those that exhibit TUR lengths > threshold). Each time, the difference of the mean half-lives of the two groups (Δ*T*_1/2_, blue, linear scale) was estimated and Mann-Whitney U-test was performed to test whether the respective distributions differ significantly. The *p*-values of the test are plotted in log scale (black). PARS score data of the last 24 nucleotides of the transcripts were inconclusive (which is why 3′ TUR length thresholds start from 25 nt, **(C)**, broken axis), though RNAfold clearly suggested that these regions are unstructured **(A)**. The plots are generated using experimental half-life data by Miller and his group (Miller *et al*, 2011). Plots for rest of the datasets are given in the supplementary (Fig S2A-B).

### RNase machineries comprise catalytic sites accessible through narrow internal tunnels

To understand the geometrical criteria of enzyme-substrate recognition associated with mRNA degradation, we collected high-resolution crystallographic structures of known RNase machines of *S. cerevisiae* (**Fig. 1C–G, Fig S1A-C**,**Table S1**) and their respective catalytic site information (Porter *et al*, 2004) (**Table S1**). A comprehensive methodology (see **Materials and Methods**) was developed to identify the degradation tunnels (with an entry and an exit pore) in these machines that single-stranded mRNAs can traverse to access the catalytic sites.

Results show that the minimum length threshold required to traverse the tunnel entry to the catalytic site are different for different RNase machineries (**Table S1**). For example, progressive 5′→3′ exoribonucleases Xrn1 and Xrn2 comprise 35–40 Å long internal tunnels that 5–6 nucleotide (nt) long single-stranded mRNA substrates can traverse (**Fig 1 C, D, Fig S1A**). The X-ray crystallographic structure of *Drosophila* Xrn1, bound to an RNA substrate (Jinek *et al*, 2011), showed very similar length requirements.

The major progressive 3′→5′ exoribonuclease, exosome (Exo-10, **Fig. 1E, Fig S1B**), is formed by nine catalytically inert subunits (Exo-9, comprising the RNase PH-like ring and the S1 and KH ring) and one catalytically active RNase, Rrp44 (Makino *et al*, 2013a, 2013b). A comprehensive RNase digestion assay investigating the lengths of single stranded RNA being protected by inactive Exo-10, against RNase A and RNase T1, detected 31–33 nt RNA fragments (Bonneau *et al*, 2009). Later, an X-ray crystallographic structure of Exo-10, bound to a substrate RNA (Makino *et al*, 2013a), confirmed that a 31–33 nt single-stranded RNA substrate can traverse the highly conserved core internal tunnel (roughly 160 Å path, Table S1) and access the catalytic site. Recently, a Cryo-EM structure of human Exo-10 (Weick *et al*, 2018), showed a very similar 31–33 nt path.

The catalytic subunit of exosome, progressive 3′→5′ exoribonuclease Rrp44, also digests unstructured substrates on its own. The previously mentioned RNase digestion assay showed that inactive Rrp44ΔPIN (Rrp44 without the endonuclease PIN domain) protects 9–10 nt RNA fragments (Bonneau *et al*, 2009). A careful analysis of Rrp44 structure unraveled an internal tunnel of ∼57 Å (Fig. 1F, Fig S1C, Table S1), that a single-stranded RNA of 9–10 nt length can traverse. X-ray and Cryo-EM structures of yeast (Bonneau *et al*, 2009) and human Exo-10 (Makino *et al*, 2013a), bound to the substrate RNAs, further confirmed this estimation. In the nucleus, the Exo-10 core recruits a distributive 3′→5′ exoribonuclease Rrp6. Rrp6 comprises a ∼49 Å internal tunnel through which 8–9 nt long substrates can traverse (Fig. 1G, Fig S1C). In summary, cells encode different classes of RNases and their geometrical criteria of substrate recognition also varies broadly.

### Substrate mRNA molecules tend to exhibit unstructured regions at both termini

A generalized feature implicated in protein degradation is that substrate proteins tend to exhibit intrinsically disordered termini (van der Lee *et al*, 2014; Lobanov *et al*, 2010; Uversky, 2013). To understand whether substrate mRNA molecules exhibit similar structural features, we combined experimental PARS-score based estimation of mRNA secondary structures (Kertesz *et al*, 2010) with RNAfold-based computational predictions (Hofacker, 2003). PARS-scores represent the single (PARS < 0) or double stranded nature (PARS > 0) of RNA molecules at single nucleotide resolution, using ribonuclease digestion and high-throughput sequencing (Wan *et al*, 2013). Based on the PARS score, the single and double stranded regions of the transcript were designated as unstructured and structured regions, respectively. RNAfold predicts the minimum free energy structure (Δ*G*) of a given RNA sequence using Zuker and Stiegler algorithm (Zuker & Stiegler, 1981). A sliding-window approach was applied, in which starting from the 5′-end of the transcript, a window of 25 nucleotides (other thresholds provide very similar results) was sliding towards the 3′-end, one nucleotide at a time, and the average PARS scores of all the nucleotides within that window was compared with their RNAfold-predicted MFE. Normalizing transcript lengths within the range 1–100, the PARS and negative MFE values (−Δ*G*), averaged across all transcripts were plotted in Fig 2A. Across the transcript lengths, both parameters depicted a clear and consistent tendency that transcripts tend to be less structured at their termini than at the middle.

### Terminal unstructured regions influence mRNA half-lives on a genome scale

Since mRNAs engage with exonuclease machineries through their terminal unstructured regions (TUR) (Houseley & Tollervey, 2009; Hui *et al*, 2014), those featuring TURs long enough to traverse the respective RNase machinery internal tunnel and access the catalytic sites would presumably be degraded faster (shorter half-life) than mRNAs without this feature. One *in vitro* biochemical study (Lorentzen *et al*, 2008) on Rrp44 (requires 9 nt 3′ TUR) showed that such principles do exist: the authors incubated duplex RNA substrates with 14 nt, 7 nt, 4 nt, and 2 nt long 3′ TURs with Rrp44 and observed that while Rrp44 efficiently degraded substrates with 14 nt long 3′ TURs, those comprising 7 nt and 4 nt long 3′ TURs degraded slower and slower, while those having 2 nt long 3′ TURs were not degraded at all.

To understand the effect of TUR lengths on mRNA half-life, the 5′ and 3′ TUR lengths of each transcript were estimated from the PARS-score data (see **Materials and Methods**). For a transcript of length *L*, if *s1* and *s2* are indices of the first and last structured nucleotides having PARS score > 0, then the 5’ and 3’ terminal unstructured region (TUR) lengths were assigned as *s1-1* and *L-s2-1*, respectively. The estimated 5′ TUR lengths varied within the range 0–300 nt (mean = 7.8, median = 2.5) (Fig S2C, Data S1); 3′ TUR lengths varied within the range 25–1164 nt (mean = 37.9, median = 30.0) (Fig S2C, Data S1). The PARS score data of the last 24 nucleotides at the 3’ UTR region of the transcripts were inconclusive (PARS = 0). However, RNAfold-predicted −Δ*G* values clearly suggested that these regions tend to be unstructured (Fig 2A).

Controlled protein degradation in eukaryotic cells is mediated by a single machinery, proteasome; meaning, the same 30 aa TDR length criterion applies to all the substrates. If we classify yeast proteins into two groups depending on the TDR lengths: short (TDR ≤ 30 aa) and long (TDR > 30 aa), the latter exhibits significantly shorter half-life than the former (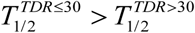, Mann Whitney U-test *p* < 10^−5^) (van der Lee *et al*, 2014). Degradation of mRNAs, on the other hand, involves multiple RNase machineries having characteristic TUR length criteria and specific directional preferences for degradation (3′→5′ or 5′→3′). Since no single length cutoff applies to all mRNAs, we varied 5′ and 3′ TUR length cutoffs, *i* and *j*, and investigated whether 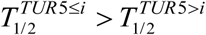 (half-lives of mRNAs having 5′ TUR ≤ *i* (short group) are longer than those having 5′ TUR > *i* (long group)) and 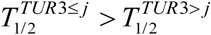 (whether half-lives of mRNAs having 3′ TUR ≤ *j* (short group) are longer than those having 3′ TUR > *j* (long group)) trends exist on a genome scale. As 5′ TUR length *i* varied, we plotted the half-life differences of short and long groups 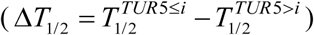 and the *p*-values representing whether 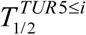 and 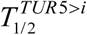 significantly differ (MWU-test of equal median).

Results showed that for different 5′ TUR thresholds, half-life difference (and the respective *p*-value) between short and long groups attains a maxima (minima) within the range 3 ≤ *i* ≤ 8 nt, and then gradually decreases (increases) as longer length thresholds were applied (Fig 2B, Fig S2A). Given that the major progressive 5′→3′ exoribonucleases Xrn1/2 require ∼5 nt 5′ TUR to engage efficiently, this result clearly depicts that mRNAs comprising 5′ TURs amenable to efficient Xrn1/2 engagement have significantly shorter half-lives than mRNAs without this feature. As longer and longer length thresholds were applied, slowly and rapidly degrading mRNAs mixed together in the short group, and the significant difference between the short and the long group continued to fade away. In case of 3′ TUR, the half-life difference (and the respective *p*-value) between short and long groups attained a maxima (minima), within the range 29 ≤ *j* ≤ 41nt, and then gradually decreased (increased) as longer length thresholds were applied (Fig 2C, Fig S2B). This 29–41 nt range probably reflects the 31-33 nt mRNA path through the Exo-10 central tunnel (Fig 1E), which extends to 40–41 nt when Exo-10 recruits ATP-dependent RNA helicase Ski-complex (cytoplasm) and TRAMP complex (nucleus, Table S1) in order to mechanically unwind structured mRNA substrates prior to degradation (Makino *et al*, 2013b). These results suggest that the geometrical criteria of enzyme-substrate recognition can be efficiently captured by analyzing high-resolution crystallographic structures of the degradation machineries and these criteria influence half-lives of the substrate molecules on a genome-scale.

### The proportion of internal structured and unstructured segments likely influence engagement to endonuclease machineries and thereby regulate transcript half-lives

Yeast genome encodes various endonuclease machineries with a wide variety of functions (reviewed in (Tomecki & Dziembowski, 2010)), among which here we focus on Rrp44-mediated controlled mRNA degradation. The N-terminal PIN domain of Rrp44 harbors endonuclease activity, that cuts substrate mRNAs from the middle. Endonuclease activity does not involve an internal tunnel (Bonneau *et al*, 2009). A careful survey of Rrp44 crystal structure showed that the key catalytic residue Asp171 is located roughly at the middle of the PIN domain surface (Lebreton *et al*, 2008; Bonneau *et al*, 2009), about 30 Å distant from the boundary of the PIN domain. Thus, one can speculate that a single stranded substrate mRNA of 8–12 nt length should be able to position itself around Asp171 (Schaeffer *et al*, 2009; Bonneau *et al*, 2009), along the PIN domain structure. Thus, using different thresholds between 8–14 nt, we tried to understand whether and to what extent the geometrical criterion for endonuclease engagement influences transcript half-lives on a genome scale.

We first developed a methodology to systematically capture the internal unstructured regions (IURs) of transcripts from experimentally determined PARS-scores (Kertesz *et al*, 2010). We adopted a sliding window approach. Starting from the 5′-end of the transcript, a window of 12 nucleotides was sliding towards the 3′-end, 6 nucleotides slide at a time (12/6). We counted the number of windows in which ≥ 60% nucleotides with PARS score < 0 (*n*_*us*_, putative unstructured regions, amenable to endonuclease engagement). The number of windows that did not satisfy this criterion were assigned as putative structured regions (*n*_*s*_, not amenable to endonuclease engagement). We estimated the overall (un)structured nature of the mRNA as *ξ* = (*n*_*us*_ − *n*_*s*_) (*n*_*us*_ + *n*_*s*_), which ranges between −1 to +1 (Data S1). The mRNAs with higher *ξ* (*i.e.*, majorly unstructured) are presumably more amenable to Rrp44 engagement, thus would have shorter half-lives. All mRNAs in our dataset comprises at least one IUR. To understand how these segments influence mRNA half-life, we classified *S. cerevisiae* mRNAs into multiple groups based on their *ξ* values. Half-life distributions of these groups differed significantly, in a way that that mRNAs having higher proportions of unstructured elements had shorter median half-lives (*p* < 10^−226^, multi-sample Kruskal Wallis test for equal median) (Fig 3A, Fig S3A). Assigning structured and unstructured regions of different window sizes (8/4, 10/5, 14/7, and 16/8) and for different nucleotide thresholds (40%, 60%, 70% nucleotides having PARS > 0) did not alter this trend (Fig S3C-G). An example plot for 12/6 window and for 40% nucleotide threshold (a window was assigned as an unstructured region if ≥40% nucleotides had PARS score < 0) is shown in Fig 3B (Fig S3B).

**Fig 3.**
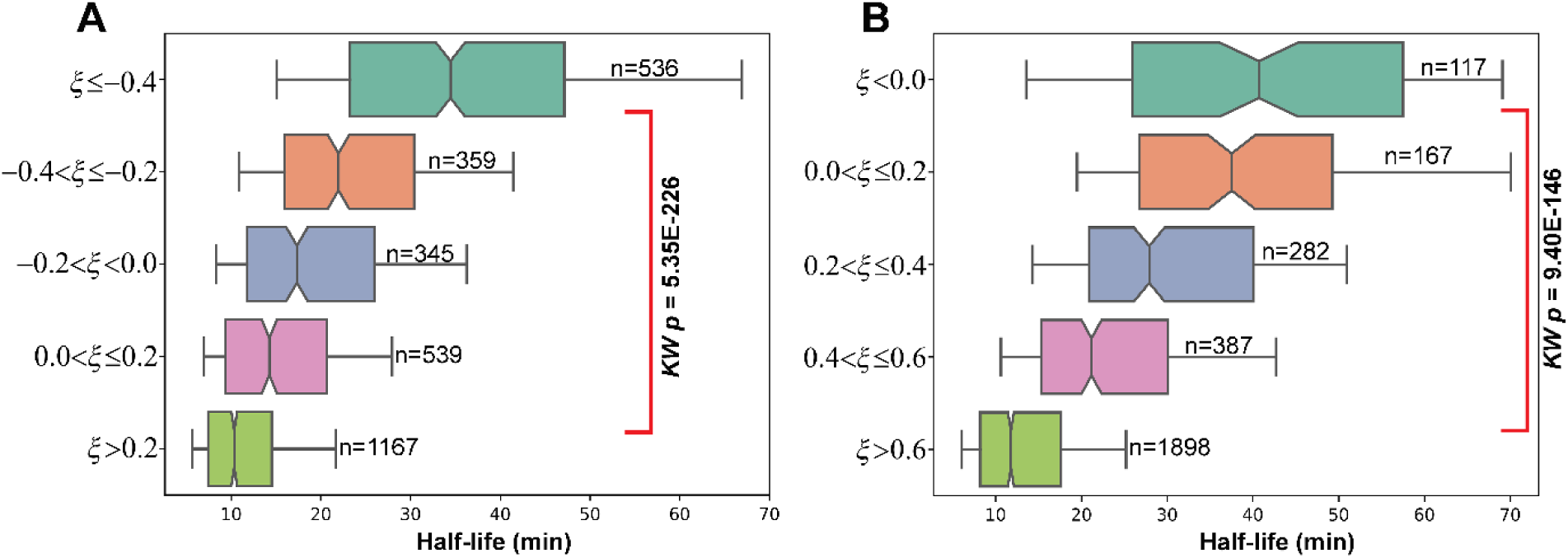
The effects of internal (un)structured regions on mRNA half-life. Half-life distributions of *S. cerevisiae* mRNAs are plotted as notched boxes (notches represent median), for different *ξ* ranges; *ξ* calculation was performed for 12/6 window size and an unstructured window was assigned for **(A)** ≥ 60% and **(B)** ≥ 40% nucleotides in the window having PARS < 0. Increasing (decreasing) *ξ* values reflect comparatively more unstructured (structured) nature of the transcript that tend to exhibit shorter (longer) half-life. Multi-sample Kruskal-Wallis test of equal median were performed to test whether the half-life distributions differ significantly, the *p*-values are provided. The plots are generated using experimental half-life data by Miller and his group (Miller *et al*, 2011). Plots for rest of the datasets are given in the supplementary (Fig S3A-B).

Overall, these results reflect that relative abundance of unstructured and structured segments within the transcripts, in which the former is a potential endonuclease engagement site and the latter is not, influences mRNA half-life on a genome scale. Previously, van der Lee *et al*, 2014 showed similar results for proteins.

### Sequestration into ribonucleoprotein complexes probably hinders exo- and endonuclease engagement and elongates transcript half-lives

Previously, working on the structural constraints influencing protein half-life on a genome-scale, we showed that sequestration into multi-component complexes warrants longer half-lives of protein subunits, presumably because the intrinsically disordered and ubiquitinoylation sites amenable to proteasomal engagement are buried under oligomeric interfaces (Mallik & Kundu, 2018). We asked whether similar molecular principles apply to mRNA degradation as well, meaning whether transcripts sequestrating into ribonucleoprotein complexes are more likely to avoid degradation (presumably by burying their unstructured regions amenable to RNase engagement at oligomeric interfaces) compared to transcripts that do not (**Fig. 4B**). This analysis was performed on a compendium of experimental protein-binding site data of *S. cerevisiae* transcripts, at single nucleotide resolution, obtained from ClipDB (Yang *et al*, 2015). This data included 50 proteins that bind to at least one of the 2665 transcripts but do not promote mRNA degradation directly or indirectly (Fig 4A, Data S2) (see **Materials and Methods**). Since transcripts can engage with RNases from both ends as well as from the internal regions, to investigate how protein binding hinders exo- and endo-nuclease activities, we mapped the protein binding sites on the transcript sequences.

**Fig 4.**
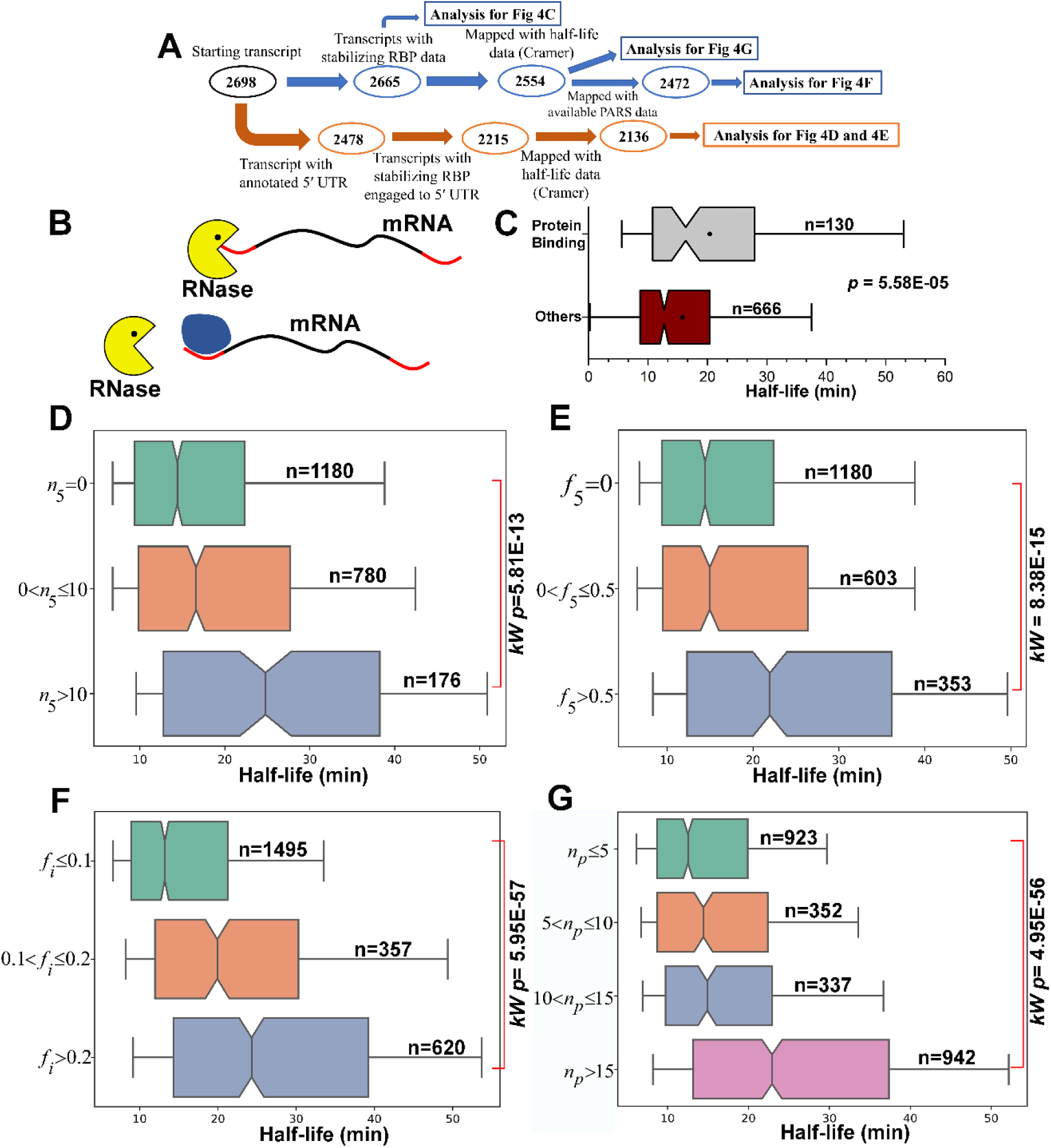
The effects of protein binding on mRNA half-life. **(A)** A sequential pipeline of the analysis along with the number of datapoints (n) used at each step **(B)** Top, a schematic representation of mRNA (black line, red termini signify the TURs) engaged with a 5′ exonuclease, represented as Pac-man; bottom, protein (blue) binding at the 5′ TUR hinders engagement. **(C)** Half-life distributions of mRNAs comprising ≥ 5 nt long 5′ TUR that do (top, grey) and do not (bottom, wine) bind to proteins. These distributions are compared by a Mann-Whitney U test, *p*-value is provided. **(D–G)** Half-life distributions of *S. cerevisiae* mRNAs (Miller et al., 2011) are plotted as notched boxes (notches represent medians), based on **(D)** the number of unique proteins (*n*_*5*_) binding to its 5′ UTR, **(E)** the fraction of the 5′ UTR length covered by protein binding regions (*f*_5_), **(F)** the fraction of internal unstructured regions that are protein binding sites (*f*_*i*_), and **(G)** the total number of unique proteins (*n*_*p*_) binding. In each case, the number of mRNAs in each group are mentioned. Multi-sample Kruskal-Wallis test of equal median was performed to test whether the distributions differ significantly; *p*-values are provided. The plots are generated using experimental half-life data by Miller and his group (Miller *et al*, 2011). The plots for rest of the datasets are given in the supplementary (Fig S4A-E).

#### Protein binding at transcript termini results in longer transcript half-life

In *S. cerevisiae*, 5′-end decapping is usually followed by rapid progressive 5′→3′ degradation by Xrn1/2 (Beelman *et al*, 1996; Muhlrad *et al*, 1994). We began with a set of 796 mRNAs comprising ≥ 5 nt long 5′ TUR and classified them into two groups: (i) those that bind to at least one protein at the 5′ TUR and (ii) those that do not. Because protein binding at 5′ TUR would, in principle, hinder Xrn1/2 engagement, the former is expected to exhibit longer half-lives than the latter. This trend was indeed observed (Fig. 4C, Fig S4A), and is statistically significant with *p* < 10^−5^ (MWU test) on a genome-scale. Next, because binding of any protein to the 5′ UTR would, in principle, hinder progressive 5′→3′ degradation, we performed the following analysis. For each transcript, (i) the number of unique proteins (*n*_*5*_) binding to its 5′ UTR (Data S2), and (ii) the fraction of the 5′ UTR length covered by protein binding regions (*f*_5_), were computed. For a given transcript, both parameters represent the probability of progressive 5′→3′ degradation being hindered by protein binding. We performed two analyses. First, based on *n*_*5*_, *S. cerevisiae* transcripts were classified into three groups: (*i*) *n*_*5*_ = 0, (*ii*) 0 < *n*_*5*_ ≤ 10, and (*iii*) *n*_*5*_ > 10. Half-lives of these three groups differed significantly (KW *p* < 10^−9^, Fig. 4D, Fig S4B), such that *T*_*1/2*_(*i*) < *T*_*1/2*_(*ii*) < *T*_*1/2*_(*iii*). Second, based on *f*_5_, transcripts were reclassified into three groups: (*a*) *f*_5_ = 0, (*b*) 0 < *f*_5_ ≤ 0.5, and (*c*) *f*_5_ > 0.5. Half-lives of these three groups again differed significantly (KW *p* < 10^−19^, Fig. 4E, Fig S4C), such that *T*_*1/2*_(*a*) < *T*_*1/2*_(*b*) < *T*_*1/2*_(*c*). These results reflect that protein binding at the 5′ UTR tunes transcript half-lives on a genome scale, likely by hindering Xrn1/2-mediated progressive 5′→3′ degradation. A similar systematic analysis could not be performed for 3′ TUR or 3′ UTR, because only for 31 transcripts protein binding sites were mapped at 3′ TURs or 3′ UTRs.

#### Protein binding at internal unstructured regions results in longer transcript half-life

In the previous section, we proposed that 8–12 nt internal unstructured segments might efficiently engage with Rrp44 PIN domain, and showed that abundance of such unstructured segments (over structured segments) results in shorter transcript half-lives. The fraction of the unstructured regions that are also protein binding sites (*f*_*i*_) were calculated for each transcript. If at least 3 nucleotides of an unstructured region comprises a protein binding site, we assigned it as a protein binding region. The parameter, *f*_*i*_, theoretically ranges from 0 to 1; and the higher the *f*_*i*_, the higher proportion of the internal unstructured regions is buried. Transcripts were classified into three groups based on *f*_*i*_ -values: (i) *f*_*i*_ ≤ 0.1, (ii) 0.2 ≤ *f*_*i*_ < 0.1 and (iii) *f*_*i*_ > 0.2, and their half-life distributions were statistically compared. Half-lives of these three groups again differed significantly (KW *p* < 10^−39^, Fig. 4F, Fig S4D), in a way that transcript half-lives elongate as higher and higher fractions of potential endonuclease engagement sites are buried at ribonucleoprotein interfaces.

#### Sequestration into multiple ribonucleoprotein complexes results in longer half-life

Previously we showed that proteins sequestrating into multiple complexes tend to have longer half-lives than those that sequestrate into only one complex, presumably because in the former case, the protein is rarely available to the proteasome machinery in its monomeric state (Mallik & Kundu, 2018). Fig. 4D indicates that similar molecular principles apply to mRNA degradation as well (transcripts that bind to multiple proteins at its 5′ UTR exhibit longer half-lives), even though, on average, 5′ UTR comprise only 28% of all the proteins binding to a transcript. Therefore, finally we classified the *S. cerevisiae* transcripts into three classes of roughly equal size, based on the number of unique proteins (*n*_*p*_) it binds to (Data S2): (*1*) *n*_*p*_ ≤ 5, (*2*) 5 < *n*_*p*_ ≤ 10, (*3*) 10 < *n*_*p*_ ≤ 15 and (*4*) *n*_*p*_ > 15. Half-lives of these four groups again differed significantly (KW *p* < 10^−56^, Fig. 4G, Fig S4E), such that *T*_*1/2*_(*1*) < *T*_*1/2*_(*2*) < *T*_*1/2*_(*3*) < *T*_*1/2*_(*4*).

These results suggest that sequestration into ribonucleoprotein complexes elevates transcript half-lives, presumably by burying the RNase engagement sites under complex interfaces. Further, promiscuous sequestration into multiple ribonucleoprotein complexes elevates transcript half-lives on a genome scale, likely because the transcript is rarely available to RNase machineries in the monomeric state.

### Differential half-lives of paralogous transcripts depend on their altered TUR lengths, modified proportions of internal unstructured segments and different oligomerization modes

Previous genome-wide analyses of yeast paralogous protein pairs (that arose from gene duplication and evolving under similar conditions), unraveled that terminal and internal disordered segments and oligomerization modes (how many macromolecular complexes they sequestrate into) of paralogous proteins diverge in the course of evolution, resulting in their altered half-lives (van der Lee *et al*, 2014; Mallik & Kundu, 2018). Based on this fact, we asked whether paralogous transcripts diverge in their 5′ and 3′ TUR lengths, in the proportions of internal structured and unstructured regions (*ξ*), and oligomerization modes (number of proteins bound to mRNA), and if they do, whether these changes correspond to their altered half-lives.

We began by classifying the paralogous pairs (Data S3) (half-lives 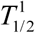 and 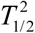) into two groups: Similar, those that during evolution maintained 5′ and 3′ TURs of roughly equal lengths (*i.e.*, 5′ TURs of both transcripts are either ≤ 5 or > 5, and 3′ TURs of both transcripts are also ≤ 33 or > 33) and Divergent, pairs with TURs of different lengths (*i.e.*, 5′ TURs of one paralog is ≤ 5, while the other is > 5, and 3′ TURs of one paralog is ≤ 33, while the other is > 33). If changes in TUR lengths correspond to changes in half-life, the ‘Divergent’ group is expected to have wider distribution of half-life differences 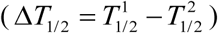 than the ‘Similar’ group. This trend was indeed observed (Fig. 5A, Fig S5A), with a *p* < 10^−6^ statistical significance (F-test under the null hypothesis of equal variances). In other words, paralogous transcripts with similar 5′ and 3′ TUR lengths tend to have similar half-lives, and TUR length dissimilarity is usually associated with differential half-lives.

**Fig 5.**
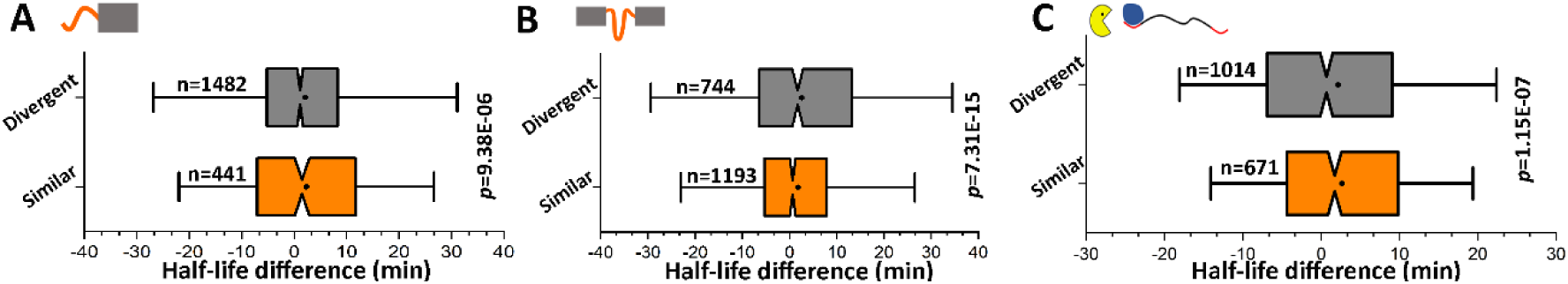
Evolutionary divergence of TUR lengths and that of internal unstructured segments influence transcript half-lives. **(A)** Distribution of half-life differences in *S. cerevisiae* paralogous transcripts, grouped according to the difference in the 5′ and 3′ TUR lengths. Similar: 5′ TURs of both transcripts are either ≤ 5 or > 5, as well as 3′ TURs are also ≤ 33 or > 33. Divergent: 5′ TUR of one paralog is ≤ 5, while the other is > 5, and 3′ TUR of one paralog is ≤ 33, while the other is > 33. **(B)** Distribution of half-life differences in *S. cerevisiae* paralogous transcripts (Miller et al., 2011), grouped according to the difference in the proportions of structured and unstructured regions [unstructured/structured: 60/40, overlap/discrete: 6/12]. Similar: −0.05 ≤ Δ*ξ* ≤ 0.05. Divergent: −0.05 > Δ*ξ* or Δ*ξ* > 0.05. **(C)** Distribution of half-life differences in *S. cerevisiae* paralogous transcripts (Miller et al., 2011), grouped according to the difference in protein binding modes. Similar: those that bind to roughly the same number of proteins, and Divergent: those that bind to differential number of proteins. For each comparison, the number of pairs in each group are mentioned, as well as the *p*-value of F-test under the null hypothesis of equal variances. The plots are generated using experimental half-life data by Miller and his group (Miller *et al*, 2011). The plots for rest of the datasets are given in the supplementary (Fig S5A-C).

A similar analysis was performed to test whether paralogous transcripts that substantially differ in the proportions of internal structured and unstructured segments (*ξ*) also show significantly larger changes in half-life than pairs that do not. For each paralogous pair, we estimated Δ*ξ* = (*ξ*_1_ − *ξ* _2_) (*ξ*_1_ + *ξ* _2_), where *ξ*_1_ and *ξ* _2_ are their proportions of internal structured and unstructured segments. These pairs were then classified into two groups: Similar, those that during evolution roughly maintained the proportions of internal structured and unstructured regions (−0.05 ≤ Δ*ξ* ≤ 0.05) and Divergent, pairs for which the proportions of internal structured and unstructured regions diverged significantly (−0.05 > Δ*ξ* or Δ*ξ* > 0.05) (Fig. 5B, Fig S5B). Half-life distributions of these two groups differed significantly (*p* < 10^−10^, F-test under the null hypothesis of equal variances) in a manner that transcript pairs with Δ*ξ* → 0 tend to exhibit highly similar half-lives and non-zero Δ*ξ* is usually associated with differential half-lives. Different Δ*ξ* thresholds did not alter our results.

We observed that oligomerization modes differ at transcript levels as well (Fig 5C, Fig S5C), which likely reflects alterations in post-transcriptional regulations after the duplication event. We specifically asked whether altered oligomerization modes lead to altered half-lives of duplicated transcripts. If two paralogous transcripts bind to *n*_1_ and *n*_2_ number of proteins, we estimated Δ*n* = (*n*_1_ − *n*_2_) (*n*_1_ + *n*_2_). Based on this parameter, paralogous transcript pairs were classified into two groups: Similar, those that bind to roughly the same number of proteins (−0.1 ≤ Δ*n* ≤ 0.1) and Divergent, those that diverged to bind differential number of proteins (−0.1 > Δ*n* or Δ*n* > 0.1). Half-life differences between these two groups differed significantly (*p* < 10^−6^, F-test under the null hypothesis of equal variances) in a manner that transcript pairs that bind to roughly the same number of proteins exhibit similar half-lives. Taken together, these results suggest that divergence of TUR lengths, of the proportions of internal structured and unstructured segments, and that of oligomerization modes generally lead to altered transcript half-lives for duplicated genes.

## Discussion

Geometrical shape complementarity is crucial to enzyme-substrate recognitions in cell. On one hand, four billion years of evolution (Hedges *et al*, 2015) has shaped both protein (Lobanov *et al*, 2010; Uversky, 2013) and mRNA molecules (Rouskin *et al*, 2014; Kertesz *et al*, 2010; Ding *et al*, 2014; Li *et al*, 2012) to exhibit intrinsically disordered/unstructured regions at their termini as well as in the middle. On the other hand, RNase and protease machineries have evolved into ‘molecular cage’-shaped nanomachines with their catalytic sites accessible through narrow internal tunnels (except endonucleases) that only these unstructured regions can traverse.

### Yeast comprises multiple RNase machineries with characteristic degradation mechanisms

In eukaryotic cells, cytoplasmic RNAs are degraded by multiple machineries. Each RNase machinery has its own geometrical criterion, and directional preference (5′→3′ or 3′→5′ or endonuclease) for substrate engagement (Houseley & Tollervey, 2009) and a characteristic mode of degradation (progressive/distributive) (Houseley & Tollervey, 2009).

A key feature implemented in mRNA degradation is that their engagement with the respective RNase machineries is the rate determining step of degradation (Laalami *et al*, 2014); this engagement involves a terminal (exonuclease) or an internal (endonuclease) unstructured region of a certain length. Despite the wide diversity of RNase machineries, PARS-score based estimation of TUR lengths showed that specific 5′ TUR and 3′ TUR length cutoffs appear to be crucial for rapid 5′→3′ and 3′→5′ degradation, and transcripts featuring TURs longer than this threshold are degraded faster than those without this feature. For 5′→3′ exonuclease degradation the aforesaid optimum 5′ TUR length for rapid 5′→3′ degradation appears to be 3–8 nt. Yeast cells comprise two related 5′→3′ exonuclease, Xrn1 and Xrn2, that require substrates with 5–6 nt long 5′ TURs for efficient engagement, which is clearly reflected in our analysis. For 3′→5′ exonuclease degradation, yeast cells comprise two machineries, Rrp6 (distributive degradation) and Rrp44 (progressive degradation). Both these enzymes can work alone, or as a part of a massive multicomponent RNase machine, the exosome. While working alone, both Rrp6 and Rrp44 can efficiently degrade short, unstructured substrate mRNAs (Lorentzen *et al*, 2008; Callahan & Butler, 2008) with ∼9 nt 3′ TUR lengths (Fig 1). However, the 3′ TUR length cutoff optimum for rapid 3′→5′ exonuclease degradation appears to be 29–41 nt. The only machinery that comprises mRNA paths of this range is the exosome. The central tunnel of Exo-10 (catalytically inert Exo-9 bound to Rrp44) incorporates a 31–33 nt mRNA path, that extends to 40–41 nt when Exo-10 binds to ATP-dependent RNA helicase Ski-complex in cytoplasm or TRAMP complex in nucleus (Table S1). Notably, these helicase complexes mechanically unwind structured mRNA substrates prior to degradation(Makino *et al*, 2013). This suggests that even though Rrp6 and Rrp44 can degrade short, unstructured RNA fragments independently (Lorentzen *et al*, 2008; Callahan & Butler, 2008), degradation of full-length yeast mRNAs, that tend to be structured (Rouskin *et al*, 2014; Kertesz *et al*, 2010; Ding *et al*, 2014; Li *et al*, 2012), seems to be predominantly mediated by the exosome itself.

The eukaryotic mRNA degradation scenario is very similar to that of protein degradation in the sense that the latter involves only one-barrel shaped machinery, Proteasome, that can engage with its substrate from either termini as well as from the middle (Ciechanover, 2005; Bhattacharyya *et al*, 2014; van der Lee *et al*, 2014). The geometrical criterion of proteasomal engagement applies to all its substrates, and proteins exhibiting disordered regions longer than these thresholds tend to have shorter half-lives than proteins without these features (van der Lee *et al*, 2014).

### Endonuclease degradation also regulates mRNA half-life on a genome-scale

Endonuclease cleavage is one of the major mRNA degradation mechanisms in the eukaryotic cell, with the resulting fragments rapidly cleared by exonuclease digestion(Schoenberg, 2011; Abernathy *et al*, 2015). The N-terminal PIN domain of Rrp44 exhibits endonuclease activity (Lorentzen *et al*, 2008). The geometrical criterion of substrate recruitment to the Rrp44 PIN domain (Rrp44PIN) and to what extent Rrp44PIN mediated endonuclease degradation influences mRNA half-life on a genome scale, have remained elusive and to our knowledge, receives its first genome-scale assessment in this study. We depicted that increasing abundance of internal unstructured segments that are presumably amenable to Rrp44PIN engagement, results in shorter transcript half-lives (Fig 3). These genome-scale trends are very similar to proteins comprising multiple ≥ 40 aa long internal disordered regions (amenable to proteasome engagement) having shorter half-lives than proteins without this feature (van der Lee *et al*, 2014).

### Sequestration into ribonucleoprotein complexes protects transcripts from degradation

Sequestration into ribonucleoprotein complexes was experimentally shown to hinder degradation of specific mRNAs in the past. Several RNA-binding proteins (HuR, for example) mask U-rich and AU-rich elements (known to promote mRNA degradation) and elevate the transcript half-life (Abdelmohsen & Gorospe,2010; Ross, 1995). Poly-(A) binding proteins can couple poly-(A) tail with the cap-binding complex, forming a loop that resists exonuclease degradation (Wakiyama *et al*, 2000; Ross, 1995). Some iron regulatory proteins are known to shield specific RNA from degradation (Ross, 1995). Despite all such case studies, to our knowledge, the effect of protein binding on mRNA stability receives its first genome-scale assessment and further exploration in this study. Working on a set of 50 *S. cerevisiae* RBPs that are not reported to promote mRNA degradation either directly or indirectly, we show that sequestration into ribonucleoprotein complexes elevates transcript half-lives across the genome and in evolution. A careful analysis of protein binding sites further suggested that oligomerization elongates transcript half-life by burying the putative exo- and endo-nuclease engagement sites (pExEnEg, terminal and internal unstructured segments) under complex interfaces, because mRNAs with larger fractions of buried pExEnEg sites tend to have longer half-life. This is very similar to oligomerization resulting in longer protein half-life, by burying the ubiquitinoylation sites and the intrinsically disordered regions necessary for proteasomal engagement under the oligomeric interfaces (Mallik & Kundu, 2018). Furthermore, association with multiple complexes results in longer protein half-life, presumably because the protein is rarely available in its monomeric form to engage with the proteasome (Mallik & Kundu, 2018). A very similar mechanism exists for mRNAs, in the sense that association with multiple proteins results in longer half-life, across the genome.

Tuning mRNA half-life by protein binding is of broad biological significance, including regulation of gene expression (Dassi, 2017; Ross, 1995) and pathogenicity (Hasan *et al*, 2014; Dassi, 2017; Moore & von Lindern, 2018). Hijacking host mRNA-stabilizing proteins to protect its own transcript is one of the most fascinating pathogenic strategies of Hepatitis C (Korf *et al*, 2005) and human papillomavirus (Cumming *et al*, 2009). Association with ribosome during translation also protects the transcript from degradation (Deana & Belasco, 2005; Edri & Tuller, 2014), and mutations promoting such phenomena often result in critical diseases. The α-Thalassemia is a well-known example, that is caused by an anti-termination mutation of UAA to CAA in the α^CS^ allele, allowing translating ribosomes to proceed into the 3′ UTR and ‘mask’ the mRNA from degradation in differentiated erythrocyte precursors (Weiss & Liebhaber, 1995). In fact, aberrant alterations of transcript half-lives had been known to lead to regulated cell death, aging and a variety of diseases (Falcone & Mazzoni, 2018; Hollams *et al*, 2002).

### Evolutionary variations of transcript half-lives that emerged from gene duplication

Gene duplication of an ancestral protein often results in sub-functionalization of the ancestral function among the duplicated copies; alternatively, non-essential or redundant functions might be lost and new functions might emerge(Lynch & Conery, 2000). Functional divergence of duplicated genes is often associated with their altered gene expression at transcript level (Ganko *et al*, 2007; Hallin & Landry, 2019) and altered molecular interactions at the protein level (Dandage & Landry, 2019; Marchant *et al*, 2019). Several genetic mechanisms may generate diversity in terminal or internal unstructured segments of duplicated transcripts and in protein binding sites that would in turn result in their altered half-lives and thereby altered gene expression. These genetic mechanisms may include mutations, insertions and deletions, expansion of tandem repeats, alternative splicing, and alternative transcription start and end sites. It is fascinating that the same genetic variations can also diversify the terminal and internal disordered region lengths and alter the surface geometry at protein level, which further results in differential oligomerization modes and thereby differential half-lives of paralogous proteins (van der Lee *et al*, 2014; Mallik & Kundu, 2018). This suggests that paralogous genes harbor genetic variations that fine-tunes their regulatory schemes in all the downstream levels of central dogma (DNA→RNA→Protein). The same mechanisms should also contribute to half-life divergence among orthologous proteins/transcripts between species.

### Additional factors that influence mRNA half-life

In addition to the geometrical criteria to engage with the respective degradation machineries, and sequestration into oligomeric complexes, additional factors, such as (i) presence of (de)stabilizing sequence and structural motifs and (ii) higher order structures of individual mRNA molecules may fine tune their half-lives. Ubiquitinoylation-specific motifs (Miller *et al*, 2004) and sequence composition of unstructured termini (that confers high-affinity engagement with proteasome) were shown to influence protein half-life on a genome-scale (Fishbain *et al*, 2015). Similarly, presence and abundance of stabilizing and destabilizing sequence and structural motifs at the 5′ and 3′ UTRs strongly influence transcript half-lives across the genome (Cheng *et al*, 2017; Rabani *et al*, 2008; Geisberg *et al*, 2014). A structural motif like stem loop promotes mRNA decay through a class of mRNA deadenylation protein which specifically targets the stem loop feature to capture its substrate (Leppek *et al*, 2013). In *Leishmania*, a certain conserved sequence signature at the 3′ UTRs trigger endonuclease activity without prior transcript deadenylation (Müller *et al*, 2010).

When a globular, folded protein is degraded, 20S proteasome works with the 19S regulatory particle to mechanically unfold the protein by pulling it from one terminal (Lee *et al*, 2001). This mechanical unfolding becomes the rate-limiting step of degradation (Bard *et al*, 2019), and because the global topology of the substrate dictates its resistance against such unfolding, the topology of the folded chain influences its degradation rate (Mallik & Kundu, 2018). Interestingly, transcripts can also fold into higher order structures (Bevilacqua *et al*, 2016; Staple & Butcher, 2005; Gutell, 2013), thus making their mechanical unwinding a prerequisite for degradation. However, unravelling such principles, awaits discovery of 3D structures of several mRNAs.

### Mechanisms of protein and mRNA degradation: similarity in diversity

In summary, four billion years of evolution have evolved versatile surveillance and quality control pathways to regularly degrade damaged protein and mRNA molecules in the cell and to replace them with newly synthesized copies. But underlying this wide diversity are remarkably similar molecular principles that regulate the turnover rates of both the molecular species on a genome-scale and in evolution. In both cases, molecular cage-shaped degradation machineries have evolved. Catalytic sites of these machines are accessible through narrow internal tunnels (except endonuclease) that only unstructured regions of the substrate molecules can traverse. These geometrical constraints of enzyme-substrate recognition are further influenced by versatile biophysical constraints, including sequestration into multicomponent complexes, (de)stabilizing structural and sequence motifs, and globular structures of individual substrate molecules that must be mechanically unfolded prior to degradation. It is remarkable that this complex interplay of versatile biophysical factors at two different levels of central dogma can be efficiently captured by analyzing the experimental 3D structures of the respective RNase and protease machineries, terminal and internal unstructured regions of the substrate molecules, and their sequestration into multicomponent complexes, despite the fact that both molecular species include nearly 1,000-fold variations of half-lives, and comprise an enormous sequence, structure, and functional diversity. These findings further promise deeper understanding of post-translational control of gene expression and associated phenomena, including cell cycle, development, circadian rhythm, ageing, and virus biology.

## Materials and Methods

### Dataset

#### S. cerevisiae transcript half-life and sequence data

Five non-redundant, experimental mRNA half-life datasets were collected (Presnyak *et al*, 2015; Miller *et al*, 2011; Neymotin *et al*, 2014; Wang *et al*, 2002; Munchel *et al*, 2011). Protein half-life data was collected from ref. (van der Lee *et al*, 2014). *S. cerevisiae* transcriptome sequences and their respective 5′ UTR, CDS and 3′ UTR annotations were obtained from Saccharomyces Genome Database (Cherry *et al*, 2012) (**Data S1**).

#### PARS data

PARS-scores represent the single (PARS score ≤ 0) or double stranded nature (PARS score > 0) of a given RNA at single nucleotide resolution (Wan *et al*, 2013), using nuclease digestion and high-throughput sequencing. Experimentally determined PARS scores of 3204 *S. cerevisiae* transcripts were obtained from (Kertesz *et al*, 2010). For 3000 of these transcripts, 5′ UTR, 3′ UTR and CDS annotations are available (**Data S1**).

#### RNA-Protein Binding data

RNA-protein binding data for *S. cerevisiae* was obtained from ClipDb (Yang *et al*, 2015). This database includes transcriptome-wide binding sites of RBPs at the single-nucleotide level identified by crosslinking immunoprecipitation experiments. We used the protein-binding sites predicted by the statistically robust, pliant computational pipeline of Piranha (Uren *et al*, 2012) for our work. This data included 61 proteins binding to 2698 transcripts (**Data S2**). These proteins were manually reviewed and 11 proteins directly or indirectly involved in mRNA degradation were removed from further analysis. Our final workable dataset includes 50 stabilizing RNA-binding protein engagement mapped to 2665 transcripts (Fig 4A, Data S2).

#### Structural data of RNase machines

From the seminal review of Houseley and Tollervey (Houseley & Tollervey, 2009), an initial list of yeast RNases was prepared. This list included 5′-exoribonucleases Xrn1 and Xrn2, 3′-exoribonucleases exosome, Rrp44, Rrp6 and exosome, and endonuclease Rrp44. High-resolution (< 3Å) crystal structures of these machineries are collected from Protein Data Bank (Berman *et al*, 2007). We also used the model structure of Xrn1 available in ModBase (Pieper *et al*, 2011). Catalytic site residues of these RNases were collected from Catalytic Site Atlas (Porter *et al*, 2004) and by literature search (**Table S1**). In addition, a crystal structure of yeast cytoplasmic Ski complex (Halbach *et al*, 2013) and a human exosome structure with nuclear MTR4 and substrate RNA were collected (Weick *et al*, 2018).

### Tunnel analysis of RNase machines

For each machinery (i) all probable tunnels having a bottleneck radius of ≥ 3.4 Å were identified in the crystal structures using Caver Analyst v2.0 (Jurcik *et al*, 2018), (ii) tunnels that pass through the catalytic sites were filtered manually, (iii) finally, uninterrupted sliding of a train of ellipsoids of 3.4 Å major and 2.9 Å minor axis lengths (approximating sliding nucleotides) from one opening of the tunnel to the other was confirmed. This analysis extracted the probable degradation tunnels in each RNase machine and further allowed us to compute the minimum lengths of unstructured single-stranded RNA required to traverse the distance from the tunnel opening to the catalytic site (**Table S1**).

### Transcript structure analysis

#### PARS-score based Terminal Unstructured Region length assignment

For a transcript of length *L*, if *s1* and *s2* are the indices of the first and last nucleotides with PARS scores > 0, then the 5′ and 3′ terminal unstructured regions (TUR) lengths were assigned as *s1*−*1* and *L*−*s2*−*1*.

#### RNAfold-based secondary structure prediction

The secondary structures of transcript sequences and associated folding free energy were predicted using RNAfold program from ViennaRNA package (Hofacker, 2003). A sliding-window approach was adopted. Starting from the 5′-end of the transcript, a window of 25 nucleotides (other thresholds rendered similar results) was sliding towards the 3′-end, one nucleotide at a time.

### Paralogue data

Paralogous transcript pairs of *S. cerevisiae* were identified based on their sequence similarities and identical domain contents at protein level. First, NCBI standalone protein-protein BLAST (Camacho *et al*, 2009) was run across the proteomes and BLAST hits were filtered with 10^−10^ e-value thresholds. A subset of these filtered pairs, that further exhibit identical domain content (domain assignments with *p*-value ≤ 10^−5^) as annotated in Pfam (Finn *et al*, 2014), were considered as paralogous pairs (**Data S3**).

### Statistical Analysis and Data visualization

All the statistical test mentioned in the main text are performed with our in-house python scripts and Past v3.0 (Hammer *et al*, 2001). All the images are produced using Pymol, OriginPro, Matplotlib and Seaborn package of Python 2.7, and Adobe Photoshop.

## Data Availability Statement

All the raw data that support the plots within this paper and other finding of this study are available as Supplementary Files.

## Acknowledgement

The authors acknowledge Dr. Devin Trudeau, Dr. Jayanta Mukhopadhyay and Dr. Runa Sur for their valuable comments.

## Author Contribution

S.M., S.B., and S.K. designed research and implemented computational methodologies, S.B., S.M., S.H., and S.K. performed research and analyzed data; S.B., S.M., S.H., and S.K. wrote the paper.

## Funding

S.B. is supported by the Center of Excellence in Systems Biology and Biomedical Engineering (TEQIP Phase III), University of Calcutta, India. S.M. is supported by the PBC-VATAT Postdoctoral Fellowship, provided by the Council for Higher Education, Israel. S.H. is supported by DBT-BINC JRF fellowship (Fellow Number: DBT-BINC/2017/CU/12).

## Notes

https://www.researchgate.net/profile/Sudipto_Basu3/publication/339310471_Genome-scale_conserved_molecular_principles_of_mRNA_half-life_regulation_Supplementary_data/data/5e4a9c81a6fdccd965ac9ae1/Genome-scale-conserved-molecular-principles-of-mRNA-half-life-regulation-Supplementary-data.zip

## References

Abdelmohsen K & Gorospe M (2010) Posttranscriptional regulation of cancer traits by HuR. Wiley Interdiscip. Rev. RNA 1: 214–29 Available at: http://www.ncbi.nlm.nih.gov/pubmed/21935886

Abernathy E, Gilbertson S, Alla R & Glaunsinger B (2015) Viral Nucleases Induce an mRNA Degradation-Transcription Feedback Loop in Mammalian Cells. Cell Host Microbe 18: 243–53 Available at: http://www.ncbi.nlm.nih.gov/pubmed/26211836

Bard JAM, Bashore C, Dong KC & Martin A (2019) The 26S Proteasome Utilizes a Kinetic Gateway to Prioritize Substrate Degradation. Cell 177: 286-298.e15 Available at: http://www.ncbi.nlm.nih.gov/pubmed/30929903

Beelman CA, Stevens A, Caponigro G, LaGrandeur TE, Hatfield L, Fortner DM & Parker R (1996) An essential component of the decapping enzyme required for normal rates of mRNA turnover. Nature 382: 642–6 Available at: http://www.ncbi.nlm.nih.gov/pubmed/8757137

Belle A, Tanay A, Bitincka L, Shamir R & O’Shea EK (2006) Quantification of protein half-lives in the budding yeast proteome. Proc. Natl. Acad. Sci. U. S. A. 103: 13004–9 Available at: http://www.ncbi.nlm.nih.gov/pubmed/16916930

Berman H, Henrick K, Nakamura H & Markley JL (2007) The worldwide Protein Data Bank (wwPDB): ensuring a single, uniform archive of PDB data. Nucleic Acids Res. 35: D301-3 Available at: http://www.ncbi.nlm.nih.gov/pubmed/17142228

Bevilacqua PC, Ritchey LE, S. Z & Assmann SM (2016) Genome-Wide Analysis of RNA Secondary Structure. Annu. Rev. Genet. 50: 235–266 Available at: http://www.ncbi.nlm.nih.gov/pubmed/27648642

Bhattacharyya S, Yu H, Mim C & Matouschek A (2014) Regulated protein turnover: snapshots of the proteasome in action. Nat. Rev. Mol. Cell Biol. 15: 122–33 Available at: http://www.ncbi.nlm.nih.gov/pubmed/24452470

Bonneau F, Basquin J, Ebert J, Lorentzen E & Conti E (2009) The yeast exosome functions as a macromolecular cage to channel RNA substrates for degradation. Cell 139: 547–59 Available at: http://www.ncbi.nlm.nih.gov/pubmed/19879841

Bresson SM, Hunter O V, Hunter AC & Conrad NK (2015) Canonical Poly(A) Polymerase Activity Promotes the Decay of a Wide Variety of Mammalian Nuclear RNAs. PLoS Genet. 11: e1005610.Available at: http://www.ncbi.nlm.nih.gov/pubmed/26484760

Callahan KP & Butler JS (2008) Evidence for core exosome independent function of the nuclear exoribonuclease Rrp6p. Nucleic Acids Res. 36: 6645–55 Available at: http://www.ncbi.nlm.nih.gov/pubmed/18940861

Camacho C, Coulouris G, Avagyan V, Ma N, Papadopoulos J, Bealer K & Madden TL (2009) BLAST+: architecture and applications. BMC Bioinformatics 10: 421 Available at: http://www.biomedcentral.com/1471-2105/10/421

Chan LY, Mugler CF, Heinrich S, Vallotton P & Weis K (2018) Non-invasive measurement of mRNA decay reveals translation initiation as the major determinant of mRNA stability. Elife 7: Available at: http://www.ncbi.nlm.nih.gov/pubmed/30192227

Cheng J, Maier KC, Avsec Ž, Rus P & Gagneur J (2017) Cis-regulatory elements explain most of the mRNA stability variation across genes in yeast. RNA 23: 1648–1659 Available at: http://www.ncbi.nlm.nih.gov/pubmed/28802259

Cherry JM, Hong EL, Amundsen C, Balakrishnan R, Binkley G, Chan ET, Christie KR, Costanzo MC, Dwight SS, Engel SR, Fisk DG, Hirschman JE, Hitz BC, Karra K, Krieger CJ, Miyasato SR, Nash RS, Park J, Skrzypek MS, Simison M, et al (2012) Saccharomyces Genome Database: the genomics resource of budding yeast. Nucleic Acids Res. 40: D700–5 Available at: http://www.ncbi.nlm.nih.gov/pubmed/22110037

Ciechanover A (2005) Proteolysis: from the lysosome to ubiquitin and the proteasome. Nat. Rev. Mol. Cell Biol. 6: 79–87 Available at: http://www.ncbi.nlm.nih.gov/pubmed/15688069

Coller J & Parker R (2004) Eukaryotic mRNA decapping. Annu. Rev. Biochem. 73: 861–90 Available at: http://www.ncbi.nlm.nih.gov/pubmed/15189161

Cromm PM & Crews CM (2017) Targeted Protein Degradation: from Chemical Biology to Drug Discovery. Cell Chem. Biol. 24: 1181–1190 Available at: http://www.ncbi.nlm.nih.gov/pubmed/28648379

Cumming SA, Chuen-Im T, Zhang J & Graham S V (2009) The RNA stability regulator HuR regulates L1 protein expression in vivo in differentiating cervical epithelial cells. Virology 383: 142–9 Available at: http://www.ncbi.nlm.nih.gov/pubmed/18986664

Dandage R & Landry CR (2019) Paralog dependency indirectly affects the robustness of human cells. Mol. Syst. Biol. 15: e8871 Available at: http://www.ncbi.nlm.nih.gov/pubmed/31556487

Dassi E (2017) Handshakes and Fights: The Regulatory Interplay of RNA-Binding Proteins. Front. Mol. Biosci. 4: 67 Available at: http://www.ncbi.nlm.nih.gov/pubmed/29034245

Deana A & Belasco JG (2005) Lost in translation: the influence of ribosomes on bacterial mRNA decay. Genes Dev. 19: 2526–33 Available at: http://www.ncbi.nlm.nih.gov/pubmed/16264189

Ding Y, Tang Y, Kwok CK, Zhang Y, Bevilacqua PC & Assmann SM (2014) In vivo genome-wide profiling of RNA secondary structure reveals novel regulatory features. Nature 505: 696–700 Available at: http://www.ncbi.nlm.nih.gov/pubmed/24270811

Edri S & Tuller T (2014) Quantifying the effect of ribosomal density on mRNA stability. PLoS One 9: e102308 Available at: http://www.ncbi.nlm.nih.gov/pubmed/25020060

Eser P, Wachutka L, Maier KC, Demel C, Boroni M, Iyer S, Cramer P & Gagneur J (2016) Determinants of RNA metabolism in the Schizosaccharomyces pombe genome. Mol. Syst. Biol. 12: 857 Available at: http://www.ncbi.nlm.nih.gov/pubmed/26883383

Falcone C & Mazzoni C (2018) RNA stability and metabolism in regulated cell death, aging and diseases. FEMS Yeast Res. 18: Available at: http://www.ncbi.nlm.nih.gov/pubmed/29986027

Finley D (2009) Recognition and processing of ubiquitin-protein conjugates by the proteasome. Annu. Rev. Biochem. 78: 477–513 Available at: http://www.ncbi.nlm.nih.gov/pubmed/19489727

Finn RD, Bateman A, Clements J, Coggill P, Eberhardt RY, Eddy SR, Heger A, Hetherington K, Holm L, Mistry J, Sonnhammer ELL, Tate J & Punta M (2014) Pfam: the protein families database. Nucleic Acids Res. 42: D222–30 Available at: http://www.ncbi.nlm.nih.gov/pubmed/24288371

Fishbain S, Inobe T, Israeli E, Chavali S, Yu H, Kago G, Babu MM & Matouschek A (2015) Sequence composition of disordered regions fine-tunes protein half-life. Nat. Struct. Mol. Biol. 22: 214–21 Available at: http://www.ncbi.nlm.nih.gov/pubmed/25643324

Franks TM & Lykke-Andersen J (2008) The control of mRNA decapping and P-body formation. Mol. Cell 32: 605–15 Available at: http://www.ncbi.nlm.nih.gov/pubmed/19061636

Ganko EW, Meyers BC & Vision TJ (2007) Divergence in expression between duplicated genes in Arabidopsis. Mol. Biol. Evol. 24: 2298–309 Available at: http://www.ncbi.nlm.nih.gov/pubmed/17670808

Geisberg J V, Moqtaderi Z, Fan X, Ozsolak F & Struhl K (2014) Global analysis of mRNA isoform half-lives reveals stabilizing and destabilizing elements in yeast. Cell 156: 812–24 Available at: http://www.ncbi.nlm.nih.gov/pubmed/24529382

Geissler R & Grimson A (2016) A position-specific 3’UTR sequence that accelerates mRNA decay. RNA Biol. 13: 1075–1077 Available at: http://www.ncbi.nlm.nih.gov/pubmed/27565004

Goldberg AL & Dice JF (1974) Intracellular protein degradation in mammalian and bacterial cells. Annu. Rev. Biochem. 43: 835–69 Available at: http://www.ncbi.nlm.nih.gov/pubmed/4604628

Gutell RR (2013) Comparative Analysis of the Higher-Order Structure of RNA. In Biophysics of RNA Folding pp 11–22. New York, NY: Springer New York Available at: https://doi.org/10.1007/978-1-4614-4954-6_2

Halbach F, Reichelt P, Rode M & Conti E (2013) The yeast ski complex: crystal structure and RNA channeling to the exosome complex. Cell 154: 814–26 Available at: http://www.ncbi.nlm.nih.gov/pubmed/23953113

Hallin J & Landry CR (2019) Regulation plays a multifaceted role in the retention of gene duplicates. PLoS Biol. 17: e3000519 Available at: http://www.ncbi.nlm.nih.gov/pubmed/31756186

Hammer O, Harper DAT & Ryan PD (2001) Past: Paleontological statistics software package for education and data analysis. Palaeontologia Electronica

Hasan A, Cotobal C, Duncan CDS & Mata J (2014) Systematic analysis of the role of RNA-binding proteins in the regulation of RNA stability. PLoS Genet. 10: e1004684 Available at: http://www.ncbi.nlm.nih.gov/pubmed/25375137

Hedges SB, Marin J, Suleski M, Paymer M & Kumar S (2015) Tree of life reveals clock-like speciation and diversification. Mol. Biol. Evol. 32: 835–45 Available at: http://www.ncbi.nlm.nih.gov/pubmed/25739733

Hofacker IL (2003) Vienna RNA secondary structure server. Nucleic Acids Res. 31: 3429–31 Available at: http://www.ncbi.nlm.nih.gov/pubmed/12824340

Hollams EM, Giles KM, Thomson AM & Leedman PJ (2002) MRNA stability and the control of gene expression: implications for human disease. Neurochem. Res. 27: 957–80 Available at: http://www.ncbi.nlm.nih.gov/pubmed/12462398

Houseley J & Tollervey D (2009) The many pathways of RNA degradation. Cell 136: 763–76 Available at: http://www.ncbi.nlm.nih.gov/pubmed/19239894

Hui MP, Foley PL & Belasco JG (2014) Messenger RNA degradation in bacterial cells. Annu. Rev. Genet. 48: 537–59 Available at: http://www.ncbi.nlm.nih.gov/pubmed/25292357

Hustedt N & Durocher D (2016) The control of DNA repair by the cell cycle. Nat. Cell Biol. 19: 1–9 Available at: http://www.ncbi.nlm.nih.gov/pubmed/28008184

Jinek M, Coyle SM & Doudna JA (2011) Coupled 5’ nucleotide recognition and processivity in Xrn1-mediated mRNA decay. Mol. Cell 41: 600–8 Available at: http://www.ncbi.nlm.nih.gov/pubmed/21362555

Jurcik A, Bednar D, Byska J, Marques SM, Furmanova K, Daniel L, Kokkonen P, Brezovsky J, Strnad O, Stourac J, Pavelka A, Manak M, Damborsky J & Kozlikova B (2018) CAVER Analyst 2.0: analysis and visualization of channels and tunnels in protein structures and molecular dynamics trajectories. Bioinformatics 34: 3586–3588 Available at: http://www.ncbi.nlm.nih.gov/pubmed/29741570

Kertesz M, Wan Y, Mazor E, Rinn JL, Nutter RC, Chang HY & Segal E (2010) Genome-wide measurement of RNA secondary structure in yeast. Nature 467: 103–7 Available at: http://www.ncbi.nlm.nih.gov/pubmed/20811459

Korf M, Jarczak D, Beger C, Manns MP & Krüger M (2005) Inhibition of hepatitis C virus translation and subgenomic replication by siRNAs directed against highly conserved HCV sequence and cellular HCV cofactors. J. Hepatol. 43: 225–34 Available at: http://www.ncbi.nlm.nih.gov/pubmed/15964661

Laalami S, Zig L & Putzer H (2014) Initiation of mRNA decay in bacteria. Cell. Mol. Life Sci. 71: 1799–828 Available at: http://www.ncbi.nlm.nih.gov/pubmed/24064983

Lebreton A, Tomecki R, Dziembowski A & Séraphin B (2008) Endonucleolytic RNA cleavage by a eukaryotic exosome. Nature 456: 993–6 Available at: http://www.ncbi.nlm.nih.gov/pubmed/19060886

Lee C, Schwartz MP, Prakash S, Iwakura M & Matouschek A (2001) ATP-dependent proteases degrade their substrates by processively unraveling them from the degradation signal. Mol. Cell 7: 627–37 Available at: http://www.ncbi.nlm.nih.gov/pubmed/11463387

Lynch M & Conery JS (2000) The evolutionary fate and consequences of duplicate genes. Science 290: 1151–5 Available at: http://www.ncbi.nlm.nih.gov/pubmed/11073452

van der Lee R, Lang B, Kruse K, Gsponer J, Sánchez de Groot N, Huynen MA, Matouschek A, Fuxreiter M & Babu MM (2014) Intrinsically disordered segments affect protein half-life in the cell and during evolution. Cell Rep. 8: 1832–1844 Available at: http://www.ncbi.nlm.nih.gov/pubmed/25220455

Leppek K, Schott J, Reitter S, Poetz F, Hammond MC & Stoecklin G (2013) Roquin promotes constitutive mRNA decay via a conserved class of stem-loop recognition motifs. Cell 153: 869–81 Available at: http://www.ncbi.nlm.nih.gov/pubmed/23663784

Li F, Zheng Q, Vandivier LE, Willmann MR, Chen Y & Gregory BD (2012) Regulatory impact of RNA secondary structure across the Arabidopsis transcriptome. Plant Cell 24: 4346–59 Available at: http://www.ncbi.nlm.nih.gov/pubmed/23150631

Lobanov MY, Furletova EI, Bogatyreva NS, Roytberg MA & Galzitskaya O V (2010) Library of disordered patterns in 3D protein structures. PLoS Comput. Biol. 6: e1000958 Available at: http://www.ncbi.nlm.nih.gov/pubmed/20976197

Lorentzen E, Basquin J, Tomecki R, Dziembowski A & Conti E (2008) Structure of the active subunit of the yeast exosome core, Rrp44: diverse modes of substrate recruitment in the RNase II nuclease family. Mol. Cell 29: 717–28 Available at: http://www.ncbi.nlm.nih.gov/pubmed/18374646

Makino DL, Baumgärtner M & Conti E (2013a) Crystal structure of an RNA-bound 11-subunit eukaryotic exosome complex. Nature 495: 70–5 Available at: http://www.ncbi.nlm.nih.gov/pubmed/23376952

Makino DL, Halbach F & Conti E (2013b) The RNA exosome and proteasome: common principles of degradation control. Nat. Rev. Mol. Cell Biol. 14: 654–60 Available at: http://www.ncbi.nlm.nih.gov/pubmed/23989960

Mallik S & Kundu S (2018) Topology and Oligomerization of Mono- and Oligomeric Proteins Regulate Their Half-Lives in the Cell. Structure 26: 869-878.e3 Available at: http://www.ncbi.nlm.nih.gov/pubmed/29804822

Marchant A, Cisneros AF, Dubé AK, Gagnon-Arsenault I, Ascencio D, Jain H, Aubé S, Eberlein C, Evans-Yamamoto D, Yachie N & Landry CR (2019) The role of structural pleiotropy and regulatory evolution in the retention of heteromers of paralogs. Elife 8: Available at: http://www.ncbi.nlm.nih.gov/pubmed/31454312

Mathieson T, Franken H, Kosinski J, Kurzawa N, Zinn N, Sweetman G, Poeckel D, Ratnu VS, Schramm M, Becher I, Steidel M, Noh K-M, Bergamini G, Beck M, Bantscheff M & Savitski MM (2018) Systematic analysis of protein turnover in primary cells. Nat. Commun. 9: 689 Available at: http://www.ncbi.nlm.nih.gov/pubmed/29449567

Miller C, Schwalb B, Maier K, Schulz D, Dümcke S, Zacher B, Mayer A, Sydow J, Marcinowski L, Dölken L, Martin DE, Tresch A & Cramer P (2011) Dynamic transcriptome analysis measures rates of mRNA synthesis and decay in yeast. Mol. Syst. Biol. 7: 458 Available at: http://www.ncbi.nlm.nih.gov/pubmed/21206491

Miller SLH, Malotky E & O’Bryan JP (2004) Analysis of the role of ubiquitin-interacting motifs in ubiquitin binding and ubiquitylation. J. Biol. Chem. 279: 33528–37 Available at: http://www.ncbi.nlm.nih.gov/pubmed/15155768

Moore KS & von Lindern M (2018) RNA Binding Proteins and Regulation of mRNA Translation in Erythropoiesis. Front. Physiol. 9: 910 Available at: http://www.ncbi.nlm.nih.gov/pubmed/30087616

Moss Bendtsen K, Jensen MH, Krishna S & Semsey S (2015) The role of mRNA and protein stability in the function of coupled positive and negative feedback systems in eukaryotic cells. Sci. Rep. 5: 13910 Available at: http://www.ncbi.nlm.nih.gov/pubmed/26365394

Muhlrad D, Decker CJ & Parker R (1994) Deadenylation of the unstable mRNA encoded by the yeast MFA2 gene leads to decapping followed by 5’-->3’ digestion of the transcript. Genes Dev. 8: 855–66 Available at: http://www.ncbi.nlm.nih.gov/pubmed/7926773

Mullen TE & Marzluff WF (2008) Degradation of histone mRNA requires oligouridylation followed by decapping and simultaneous degradation of the mRNA both 5’ to 3’ and 3’ to 5’. Genes Dev. 22: 50–65 Available at: http://www.ncbi.nlm.nih.gov/pubmed/18172165

Müller M, Padmanabhan PK, Rochette A, Mukherjee D, Smith M, Dumas C & Papadopoulou B (2010) Rapid decay of unstable Leishmania mRNAs bearing a conserved retroposon signature 3’-UTR motif is initiated by a site-specific endonucleolytic cleavage without prior deadenylation. Nucleic Acids Res. 38: 5867–83 Available at: http://www.ncbi.nlm.nih.gov/pubmed/20453029

Munchel SE, Shultzaberger RK, Takizawa N & Weis K (2011) Dynamic profiling of mRNA turnover reveals gene-specific and system-wide regulation of mRNA decay. Mol. Biol. Cell 22: 2787–95 Available at: http://www.ncbi.nlm.nih.gov/pubmed/21680716

Narsai R, Howell KA, Millar AH, O’Toole N, Small I & Whelan J (2007) Genome-wide analysis of mRNA decay rates and their determinants in Arabidopsis thaliana. Plant Cell 19: 3418–36 Available at: http://www.ncbi.nlm.nih.gov/pubmed/18024567

Neymotin B, Athanasiadou R & Gresham D (2014) Determination of in vivo RNA kinetics using RATE-seq. RNA 20: 1645–52 Available at: http://www.ncbi.nlm.nih.gov/pubmed/25161313

Pieper U, Webb BM, Barkan DT, Schneidman-Duhovny D, Schlessinger A, Braberg H, Yang Z, Meng EC, Pettersen EF, Huang CC, Datta RS, Sampathkumar P, Madhusudhan MS, Sjölander K, Ferrin TE, Burley SK & Sali A (2011) ModBase, a database of annotated comparative protein structure models, and associated resources. Nucleic Acids Res. 39: D465–74 Available at: http://www.ncbi.nlm.nih.gov/pubmed/21097780

Porter CT, Bartlett GJ & Thornton JM (2004) The Catalytic Site Atlas: a resource of catalytic sites and residues identified in enzymes using structural data. Nucleic Acids Res. 32: D129–33 Available at: http://www.ncbi.nlm.nih.gov/pubmed/14681376

Presnyak V, Alhusaini N, Chen Y-H, Martin S, Morris N, Kline N, Olson S, Weinberg D, Baker KE, Graveley BR & Coller J (2015) Codon optimality is a major determinant of mRNA stability. Cell 160: 1111–24 Available at: http://www.ncbi.nlm.nih.gov/pubmed/25768907

Price JC, Guan S, Burlingame A, Prusiner SB & Ghaemmaghami S (2010) Analysis of proteome dynamics in the mouse brain. Proc. Natl. Acad. Sci. U. S. A. 107: 14508–13 Available at: http://www.ncbi.nlm.nih.gov/pubmed/20699386

Rabani M, Kertesz M & Segal E (2008) Computational prediction of RNA structural motifs involved in posttranscriptional regulatory processes. Proc. Natl. Acad. Sci. U. S. A. 105: 14885–90 Available at: http://www.ncbi.nlm.nih.gov/pubmed/18815376

Raghavan A, Ogilvie RL, Reilly C, Abelson ML, Raghavan S, Vasdewani J, Krathwohl M & Bohjanen PR (2002) Genome-wide analysis of mRNA decay in resting and activated primary human T lymphocytes. Nucleic Acids Res. 30: 5529–38 Available at: http://www.ncbi.nlm.nih.gov/pubmed/12490721

Ross J (1995) mRNA stability in mammalian cells. Microbiol. Rev. 59: 423–50 Available at: http://www.ncbi.nlm.nih.gov/pubmed/7565413

Rouskin S, Zubradt M, Washietl S, Kellis M & Weissman JS (2014) Genome-wide probing of RNA structure reveals active unfolding of mRNA structures in vivo. Nature 505: 701–5 Available at: http://www.ncbi.nlm.nih.gov/pubmed/24336214

Schaeffer D, Tsanova B, Barbas A, Reis FP, Dastidar EG, Sanchez-Rotunno M, Arraiano CM & van Hoof A (2009) The exosome contains domains with specific endoribonuclease, exoribonuclease and cytoplasmic mRNA decay activities. Nat. Struct. Mol. Biol. 16: 56–62 Available at: http://www.ncbi.nlm.nih.gov/pubmed/19060898

Schimke RT & Doyle D (1970) Control of enzyme levels in animal tissues. Annu. Rev. Biochem. 39: 929–76 Available at: http://www.ncbi.nlm.nih.gov/pubmed/4394639

Schneider C, Leung E, Brown J & Tollervey D (2009) The N-terminal PIN domain of the exosome subunit Rrp44 harbors endonuclease activity and tethers Rrp44 to the yeast core exosome. Nucleic Acids Res. 37: 1127–40 Available at: http://www.ncbi.nlm.nih.gov/pubmed/19879841

Schoenberg DR (2011) Mechanisms of endonuclease-mediated mRNA decay. Wiley Interdiscip. Rev. RNA 2: 582–600 Available at: http://www.ncbi.nlm.nih.gov/pubmed/21957046

Shabalina SA, Ogurtsov AY & Spiridonov NA (2006) A periodic pattern of mRNA secondary structure created by the genetic code. Nucleic Acids Res. 34: 2428–37 Available at: http://www.ncbi.nlm.nih.gov/pubmed/16682450

Slomovic S, Fremder E, Staals RHG, Pruijn GJM & Schuster G (2010) Addition of poly(A) and poly(A)-rich tails during RNA degradation in the cytoplasm of human cells. Proc. Natl. Acad. Sci. U. S. A. 107: 7407–12 Available at: http://www.ncbi.nlm.nih.gov/pubmed/20368444

Staple DW & Butcher SE (2005) Pseudoknots: RNA structures with diverse functions. PLoS Biol. 3: e213 Available at: http://www.ncbi.nlm.nih.gov/pubmed/15941360

Thrower JS, Hoffman L, Rechsteiner M & Pickart CM (2000) Recognition of the polyubiquitin proteolytic signal. EMBO J. 19: 94–102 Available at: http://www.ncbi.nlm.nih.gov/pubmed/10619848

Tomecki R & Dziembowski A (2010) Novel endoribonucleases as central players in various pathways of eukaryotic RNA metabolism. RNA 16: 1692–724 Available at: http://www.ncbi.nlm.nih.gov/pubmed/27151978

Uren PJ, Bahrami-Samani E, Burns SC, Qiao M, Karginov F V, Hodges E, Hannon GJ, Sanford JR, Penalva LOF & Smith AD (2012) Site identification in high-throughput RNA-protein interaction data. Bioinformatics 28: 3013–20 Available at: http://www.ncbi.nlm.nih.gov/pubmed/23024010

Uversky VN (2013) The most important thing is the tail: multitudinous functionalities of intrinsically disordered protein termini. FEBS Lett. 587: 1891–901 Available at: http://www.ncbi.nlm.nih.gov/pubmed/23665034

Wakiyama M, Imataka H & Sonenberg N (2000) Interaction of eIF4G with poly(A)-binding protein stimulates translation and is critical for Xenopus oocyte maturation. Curr. Biol. 10: 1147–50 Available at: http://www.ncbi.nlm.nih.gov/pubmed/10996799

Wan Y, Qu K, Ouyang Z & Chang HY (2013) Genome-wide mapping of RNA structure using nuclease digestion and high-throughput sequencing. Nat. Protoc. 8: 849–69 Available at: http://www.ncbi.nlm.nih.gov/pubmed/23558785

Wang Y, Liu CL, Storey JD, Tibshirani RJ, Herschlag D & Brown PO (2002) Precision and functional specificity in mRNA decay. Proc. Natl. Acad. Sci. U. S. A. 99: 5860–5 Available at: http://www.ncbi.nlm.nih.gov/pubmed/11972065

Weick E-M, Puno MR, Januszyk K, Zinder JC, DiMattia MA & Lima CD (2018) Helicase-Dependent RNA Decay Illuminated by a Cryo-EM Structure of a Human Nuclear RNA Exosome-MTR4 Complex. Cell 173: 1663-1677.e21 Available at: http://www.ncbi.nlm.nih.gov/pubmed/29906447

Weiss IM & Liebhaber SA (1995) Erythroid cell-specific mRNA stability elements in the alpha 2-globin 3’ nontranslated region. Mol. Cell. Biol. 15: 2457–65 Available at: http://www.ncbi.nlm.nih.gov/pubmed/7739530

West S, Gromak N, Norbury CJ & Proudfoot NJ (2006) Adenylation and exosome-mediated degradation of cotranscriptionally cleaved pre-messenger RNA in human cells. Mol. Cell 21: 437–43 Available at: http://www.ncbi.nlm.nih.gov/pubmed/16455498

Yang E, van Nimwegen E, Zavolan M, Rajewsky N, Schroeder M, Magnasco M & Darnell JE (2003) Decay rates of human mRNAs: correlation with functional characteristics and sequence attributes. Genome Res. 13: 1863–72 Available at: http://www.ncbi.nlm.nih.gov/pubmed/12902380

Yang Y-CT, Di C, Hu B, Zhou M, Liu Y, Song N, Li Y, Umetsu J & Lu ZJ (2015) CLIPdb: a CLIP-seq database for protein-RNA interactions. BMC Genomics 16: 51 Available at: http://www.ncbi.nlm.nih.gov/pubmed/25652745

Zuker M & Stiegler P (1981) Optimal computer folding of large RNA sequences using thermodynamics and auxiliary information. Nucleic Acids Res. 9: 133–48 Available at: http://www.ncbi.nlm.nih.gov/pubmed/6163133

