## Supplementary Figures for "Genome-scale conserved molecular principles of mRNA half-life regulation"

<u>**Supplementary Figure related to Figure 1 of main text**</u>

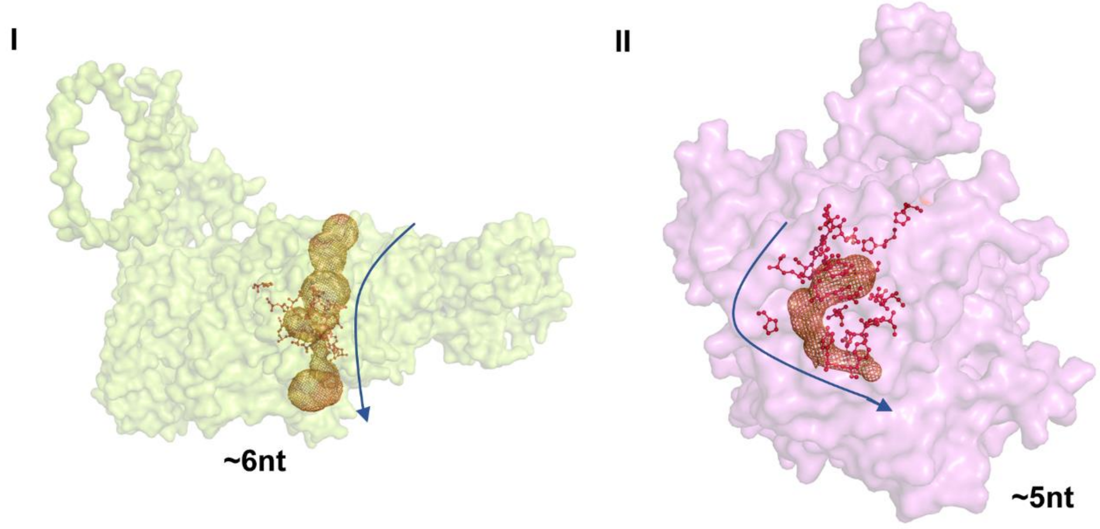


**Fig S1A. Surface representation of the 5′→3′ exoribonuclease machineries** (I) Xrn1 (modelled structure) and (II) Xrn2 (PDB: 3FQD); degradation tunnels in each machinery is represented as mesh; catalytic residues (Table S1) are shown as red sticks. Here, Xrn1 was a modelled structure, while Xrn2 is a crystallographic structure. Both the structures are modelled/crystalized without the substrate.

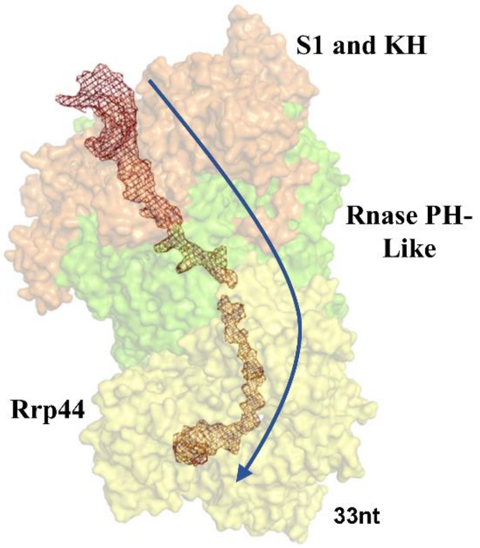


**Fig S1B.** Surface representation of the Exo-10 (comprising RNase PH-like subunits, green, and S1 and KH ring, orange) associated with progressive 3′→5′ exoribonuclease Rrp44 (yellow) (PDB: 4IFD). The structure was crystalized along with the substrate; the substrate path along the Exo-10 is illustrated as a mesh.

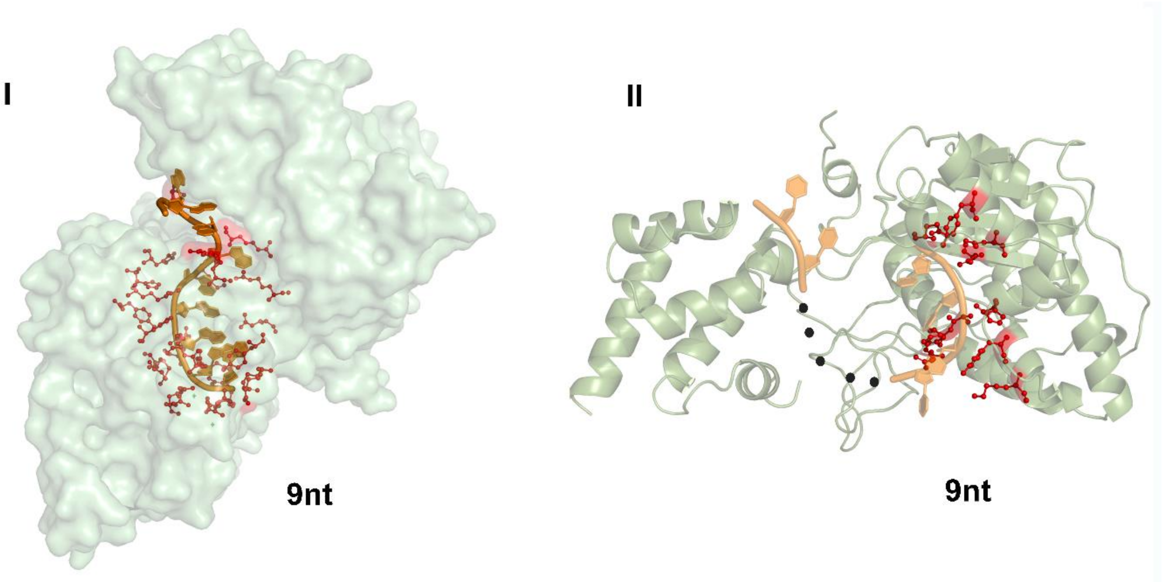


**Fig S1C.** 3′→5′ exoribonuclease machinery (I) Surface representation of the Rrp44 (PDB: 2VNU) and (II) Cartoon representation of Rrp6 (PDB: 5C0Y) along with the substrate (shown as cartoon); catalytic residues (Table S1) are shown as red sticks. In both the Rrp44 and Rrp6, the structure was resolved along with an RNA substrate. Rrp6, the substrate structure was partially resolved. The missing regions in the electron density map are highlighted by a black dotted line.

#### Supplementary Figure related to Figure 2 of main text

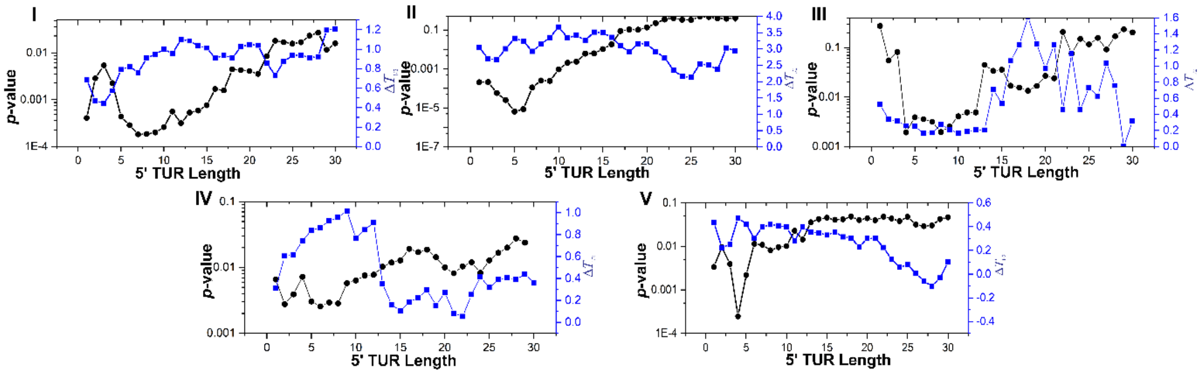


**Fig S2A.** The effects of 5′ terminal unstructured region lengths on *S. cerevisiae* mRNA half-life calculated for different datasets (I) Coller, (II) Gresham, (III) Wang1, (IV) Wang2, and (V) Weis. For different 5' TUR length threshold, yeast transcripts were classified into two groups: short (those that exhibit TUR lengths ≤ threshold) and long (those that exhibit TUR lengths > threshold). Each time, the difference of the mean half-lives of the two groups ($\Delta T_{1/2}$, blue, linear scale) was estimated and Mann-Whitney U-test was performed to test whether the respective distributions differ significantly. The *p*-values of the test are plotted in log scale (black).

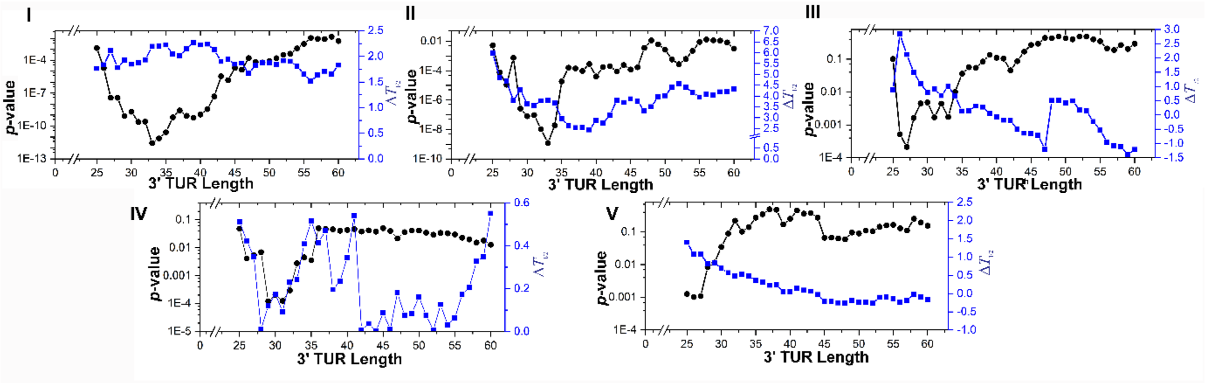


**Fig S2B.** The effects of 3′ terminal unstructured region lengths on *S. cerevisiae* mRNA half-life calculated for different datasets (I) Coller, (II) Gresham, (III) Wang1, (IV) Wang2, and (V) Weis. For different 3' TUR length threshold, yeast transcripts were classified into two groups: short (those that exhibit TUR lengths ≤ threshold) and long (those that exhibit TUR lengths > threshold). Each time, the difference of the mean half-lives of the two groups ($\Delta T_{1/2}$, blue, linear scale) was estimated and Mann-Whitney U-test was performed to test whether the respective distributions differ significantly. The *p*-values of the test are plotted in log scale (black).

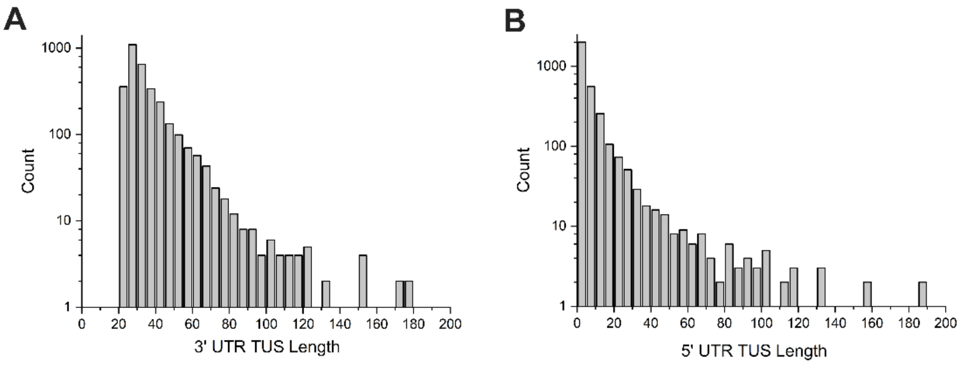


**Fig S2C.** The terminal unstructured length distribution calculated from PARS score for (A) 3′ UTR and (B) 5′ UTR. For a transcript of length *L*, if *s1* and *s2* are indices of the first and last structured nucleotides having PARS score > 0, then the 5' and 3' terminal unstructured region (TUR) lengths were assigned as *s1-1* and *L-s2-1*, respectively. The TUR is calculated based on the afore mentioned formula and their distribution is plotted. The estimated 5′ TUR lengths varied within the range 0–300 nt (mean = 7.8, median = 2.5) (Data S1); 3′ TUR lengths varied within the range 25–1164 nt (mean = 37.9, median = 30.0) (Data S1).

<u>**Supplementary Figure related to Figure 3 of main text**</u>

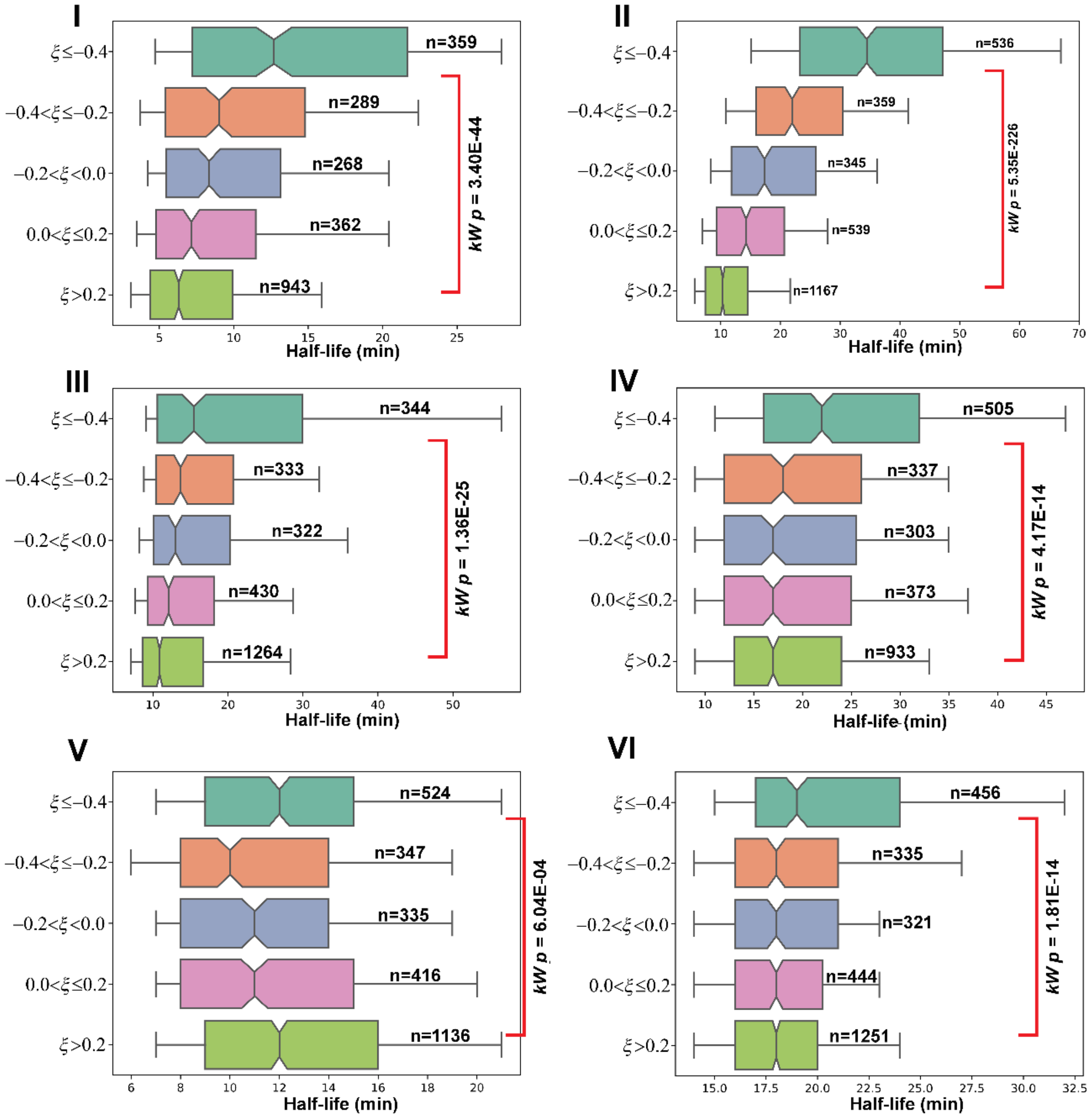


**Fig S3A.** Half-life distributions of *S. cerevisiae* mRNAs are plotted as notched boxes (notchs represent median), for different $\xi$ ranges; $\xi$ calculation was performed for 12/6 window size (i.e., a window of 12 nt was sliding from 5′ to 3′ end, 6 nt slide at a time) and an unstructured window was assigned if $\geq$ 60% nucleotides within the window have PARS $<$ 0. Increasing (decreasing) $\xi$ values reflect comparatively more unstructured (structured) nature of the transcript that tend to exhibit shorter (longer) half-life. Multi-sample Kruskal-Wallis test of equal median were performed to test whether the half-life distributions differ significantly, the *p*-values are provided. The plots are generated using experimental half-life data of (I) Coller, (II) Cramer, (III) Gresham, (IV) Wang1, (V) Wang2 and (VI) Weis datasets. Increasing (decreasing) $\xi$ values reflect comparatively more unstructured (structured) nature of the transcript that tend to exhibit shorter (longer) half-life.

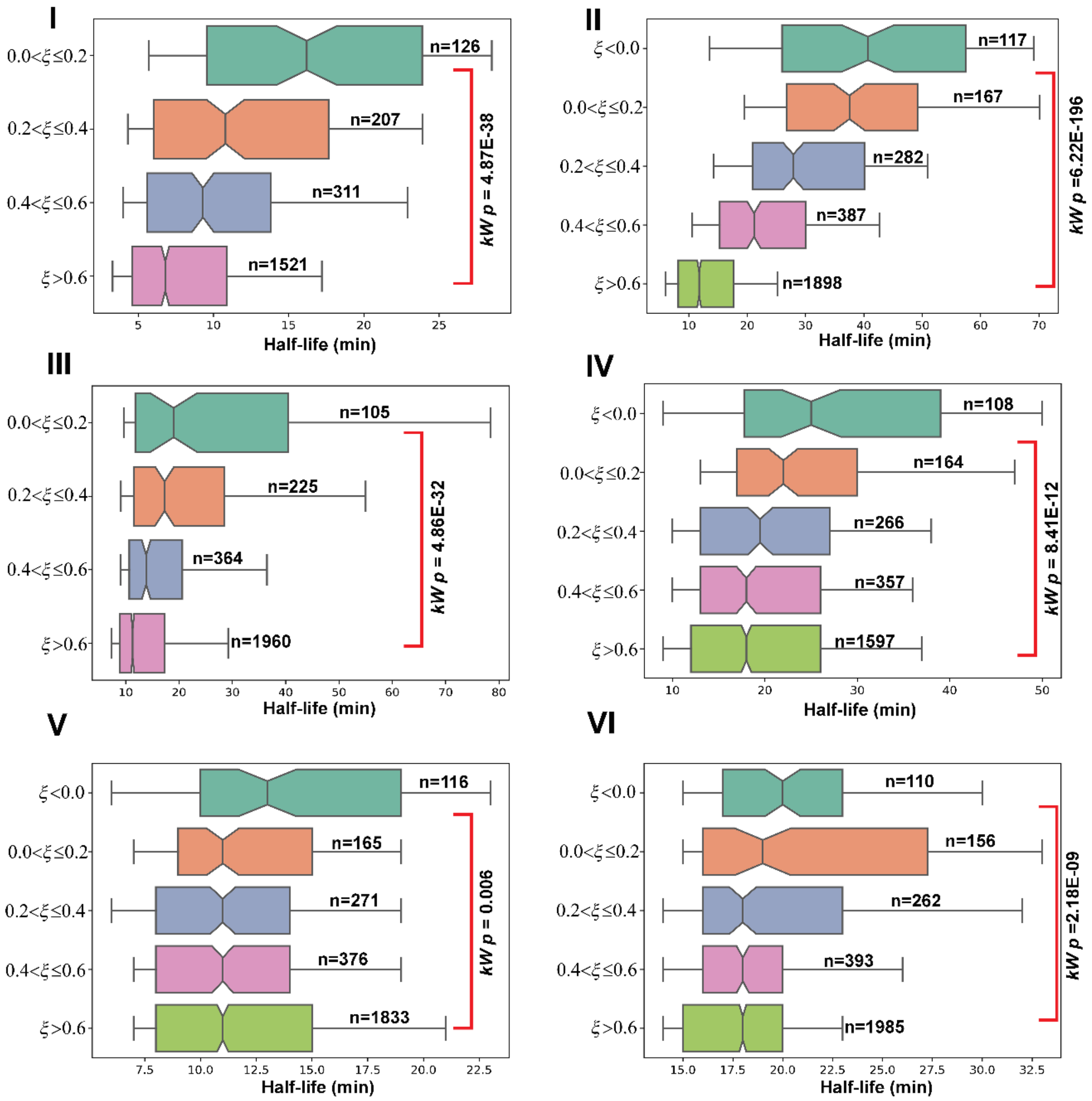


**Fig S3B.** Half-life distributions of *S. cerevisiae* mRNAs are plotted as notched boxes (notch represent median), for different $\xi$ ranges; $\xi$ calculation was performed for 12/6 window size (i.e., a window of 12 nt was sliding from 5′ to 3′ end, 6 nt slide at a time) and an unstructured window was assigned if ≥ 40% nucleotides within the window have PARS < 0. Increasing (decreasing) $\xi$ values reflect comparatively more unstructured (structured) nature of the transcript that tend to exhibit shorter (longer) half-life. Multi-sample Kruskal-Wallis test of equal median were performed to test whether the half-life distributions differ significantly, the *p*-values are provided. The plots are generated using experimental half-life data of (I) Coller, (II) Cramer, (III) Gresham, (IV) Wang1, (V) Wang2 and (VI) Weis datasets. Increasing (decreasing) $\xi$ values reflect comparatively more unstructured (structured) nature of the transcript that tend to exhibit shorter (longer) half-life.

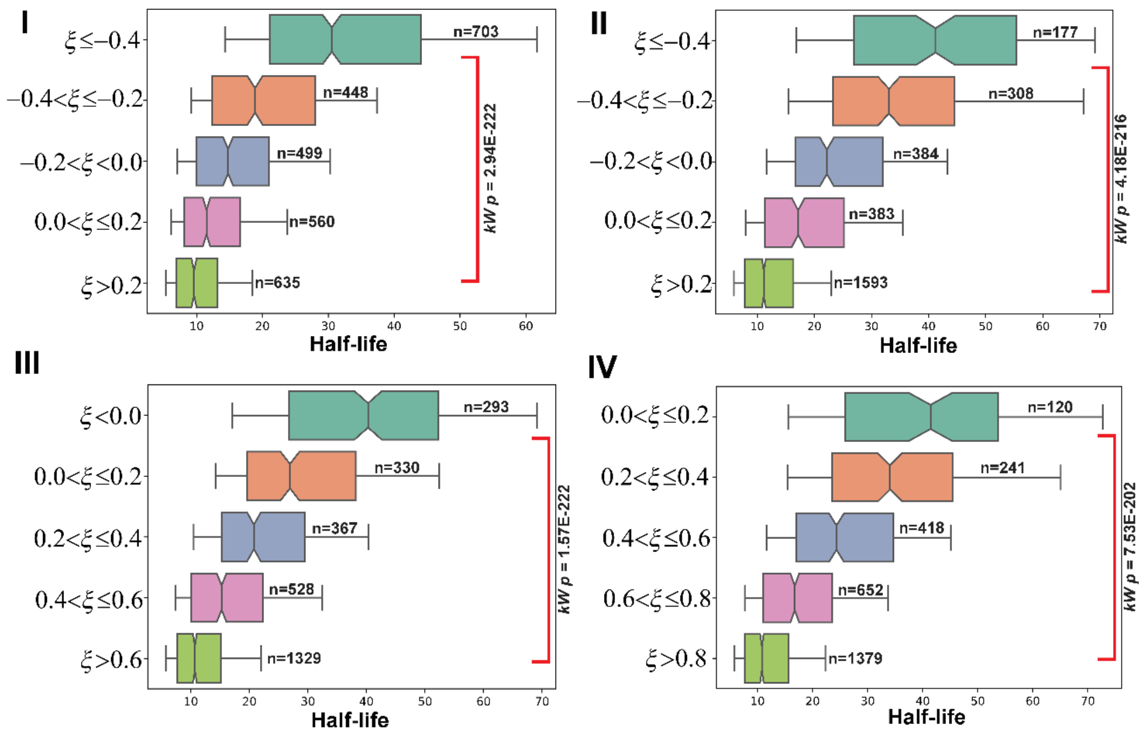


**Fig S3C.** Half-life distributions of *S. cerevisiae* mRNAs for Cramer dataset are plotted as notched boxes (notches represent median), for different $\xi$ ranges. These plots were generated using a 8/4 window (i.e., a window of 8 nt was sliding from 5′ to 3′ end, 4 nt slide at a time). A window of 8 nt was assigned as unstructured if the number of nucleotides within the window that have PARS < 0 exceeds a certain threshold. In different panels, this threshold is (I) 70% (II) 60% (III) 40% (IV) 30%. For the 8/4 window, for the said threshold, $\xi$ values were calculated; transcripts were classified into different groups according to the $\xi$-value and their half-life distributions were plotted. These distributions were compared by multi-sample Kruskal-Wallis test of equal median, the *p*-values are provided.

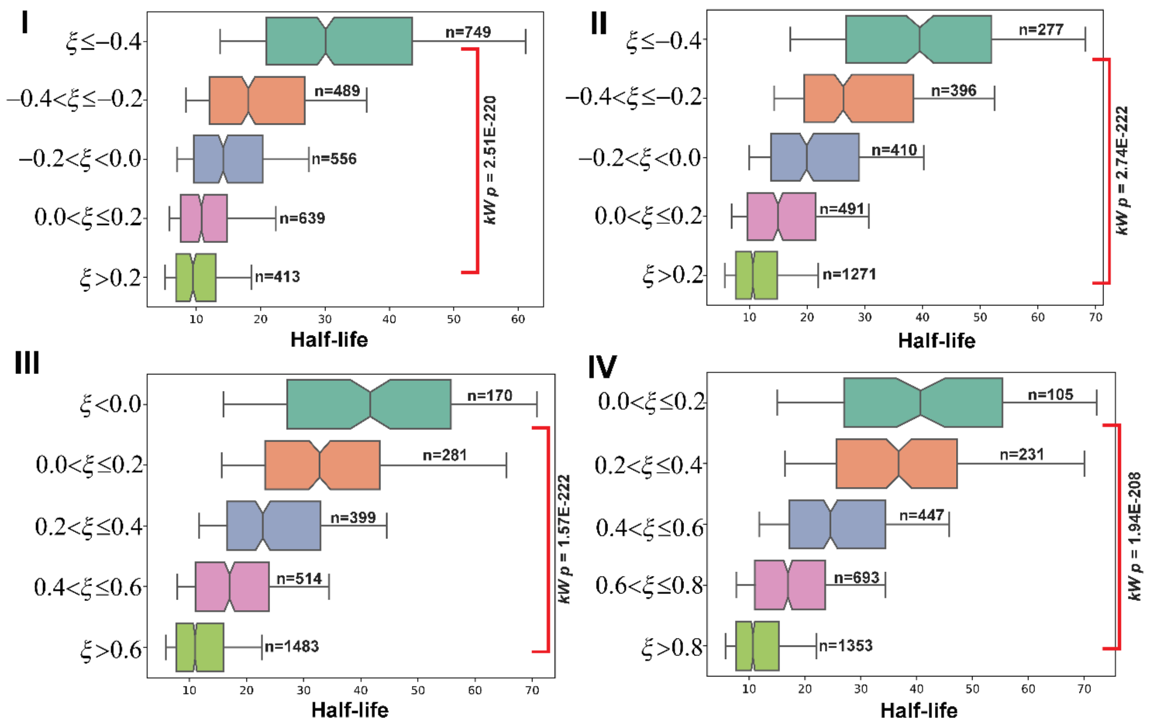


**Fig S3D.** Half-life distributions of *S. cerevisiae* mRNAs for Cramer dataset are plotted as notched boxes (notch represent median), for different $\xi$ ranges. These plots were generated using a 10/5 window (i.e., a window of 10 nt was sliding from 5′ to 3′ end, 5 nt slide at a time). A window of 10 nt was assigned as unstructured if the number of nucleotides within the window that have PARS < 0 exceeds a certain threshold. In different panels, this threshold is (I) 70% (II) 60% (III) 40% (IV) 30%. For the 8/4 window, for the said threshold, $\xi$ values were calculated; transcripts were classified into different groups according to the $\xi$-value and their half-life distributions were plotted. These distributions were compared by multi-sample Kruskal-Wallis test of equal median, the *p*-values are provided.

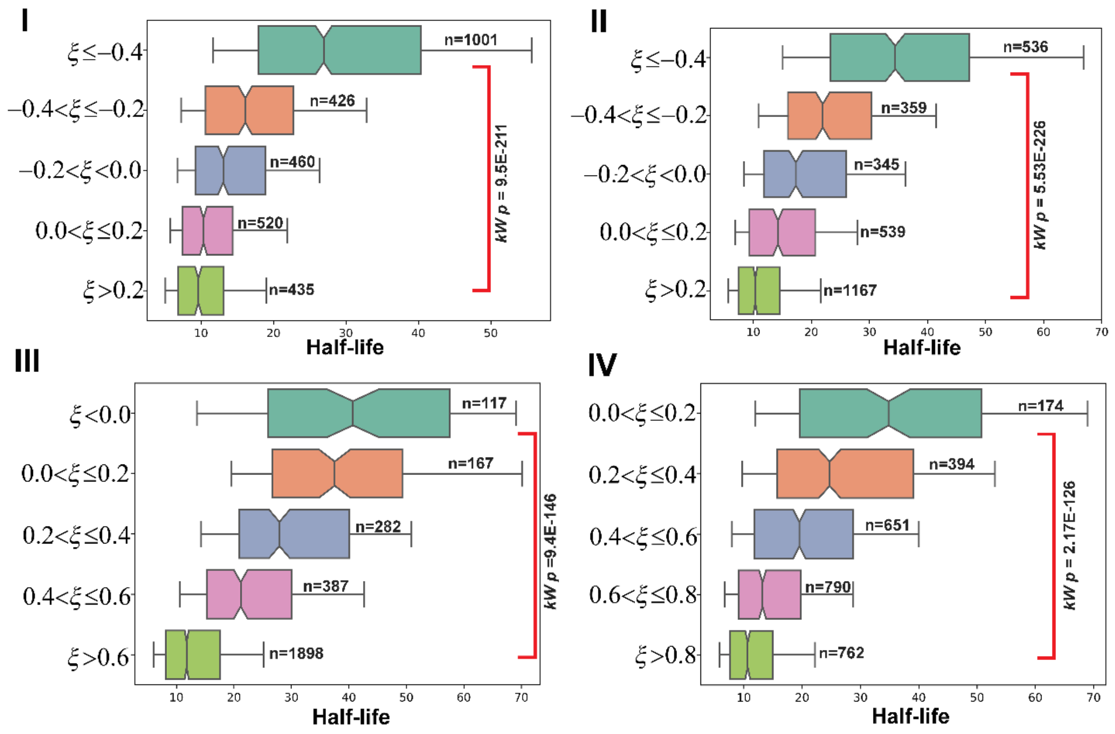


**Fig S3E.** Half-life distributions of *S. cerevisiae* mRNAs for Cramer dataset are plotted as notched boxes (notch represent median), for different $\xi$ ranges. These plots were generated using a 12/6 window (i.e., a window of 12 nt was sliding from 5′ to 3′ end, 6 nt slide at a time). A window of 12 nt was assigned as unstructured if the number of nucleotides within the window that have PARS < 0 exceeds a certain threshold. In different panels, this threshold is (I) 70% (II) 60% (III) 40% (IV) 30%. For the 8/4 window, for the said threshold, $\xi$ values were calculated; transcripts were classified into different groups according to the $\xi$-value and their half-life distributions were plotted. These distributions were compared by multi-sample Kruskal-Wallis test of equal median, the *p*-values are provided.

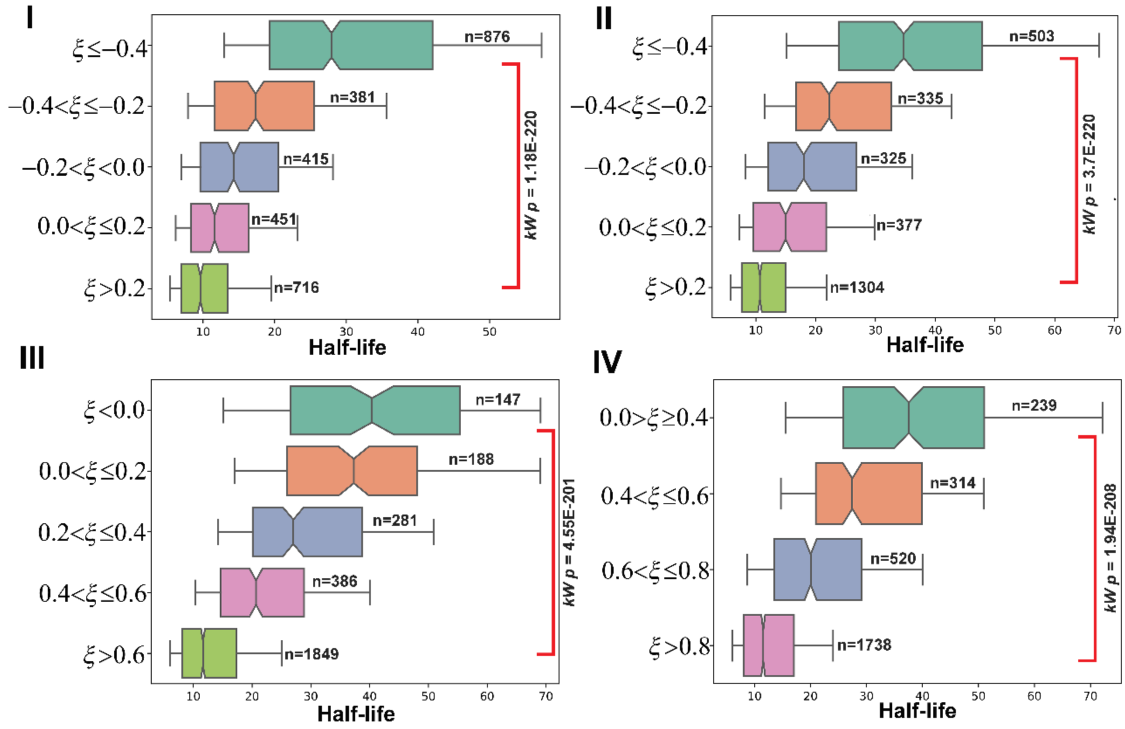


**Fig S3F.** Half-life distributions of *S. cerevisiae* mRNAs for Cramer dataset are plotted as notched boxes (notch represent median), for different $\xi$ ranges. These plots were generated using a 14/7 window (i.e., a window of 14 nt was sliding from 5′ to 3′ end, 7 nt slide at a time). A window of 14 nt was assigned as unstructured if the number of nucleotides within the window that have PARS < 0 exceeds a certain threshold. In different panels, this threshold is (I) 70% (II) 60% (III) 40% (IV) 30%. For the 8/4 window, for the said threshold, $\xi$ values were calculated; transcripts were classified into different groups according to the $\xi$-value and their half-life distributions were plotted. These distributions were compared by multi-sample Kruskal-Wallis test of equal median, the *p*-values are provided.

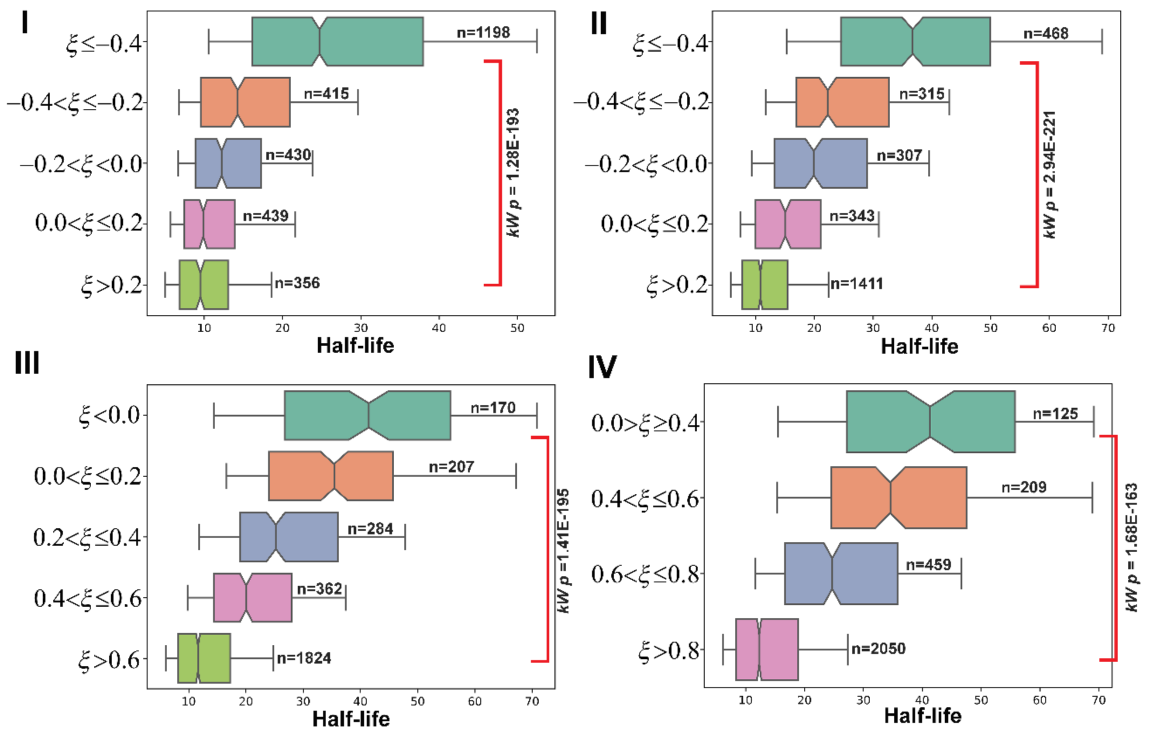


**Fig S3G.** Half-life distributions of *S. cerevisiae* mRNAs for Cramer dataset are plotted as notched boxes (notch represent median), for different $\xi$ ranges. These plots were generated using a 16/8 window (i.e., a window of 16 nt was sliding from 5′ to 3′ end, 8 nt slide at a time). A window of 16 nt was assigned as unstructured if the number of nucleotides within the window that have PARS < 0 exceeds a certain threshold. In different panels, this threshold is (I) 70% (II) 60% (III) 40% (IV) 30%. For the 8/4 window, for the said threshold, $\xi$ values were calculated; transcripts were classified into different groups according to the $\xi$-value and their half-life distributions were plotted. These distributions were compared by multi-sample Kruskal-Wallis test of equal median, the *p*-values are provided.

### Supplementary Figure related to Figure 4 of main text

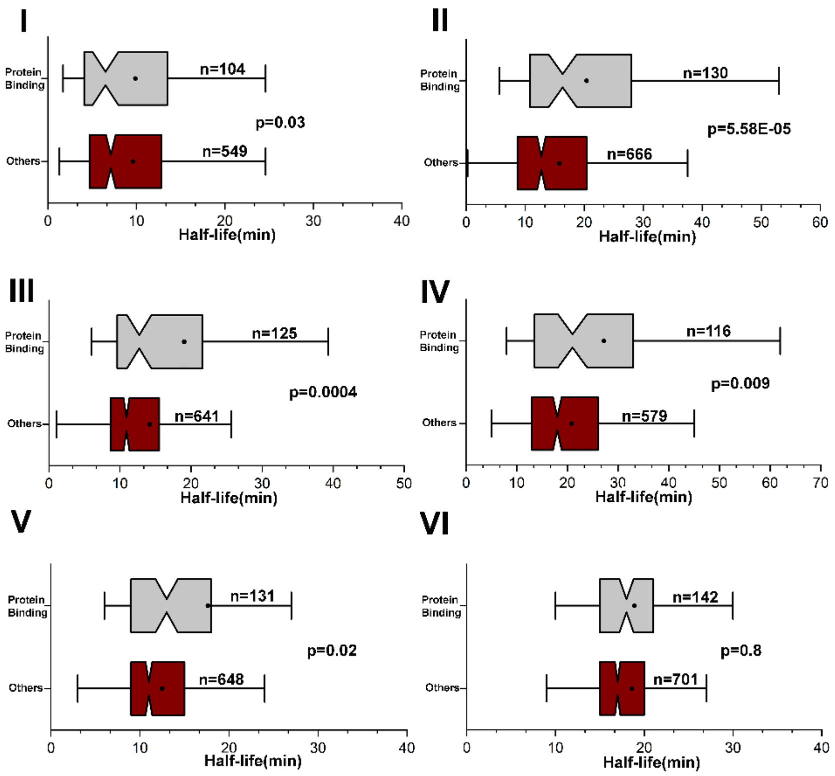


**Fig S4A.** Half-life distributions of mRNAs comprising ≥ 5 nt long 5′ TURs that do (top, grey) and do not (bottom, wine) bind to a protein, for different datasets (I) Coller, (II) Cramer, (III) Gresham, (IV) Wang1, (V) Wang2, and (VI) Weis. These distributions were compared by Mann-Whitney U tests, *p*-values are provided. 5′→3′ exonuclease machineries Xrn1 and Xrn2 require ~5 nt 5′ TUR for efficient engagement. In all these comparisons, protein binding results in longer half-lives of the respective transcripts, presumably by hindering the exonuclease digestion.

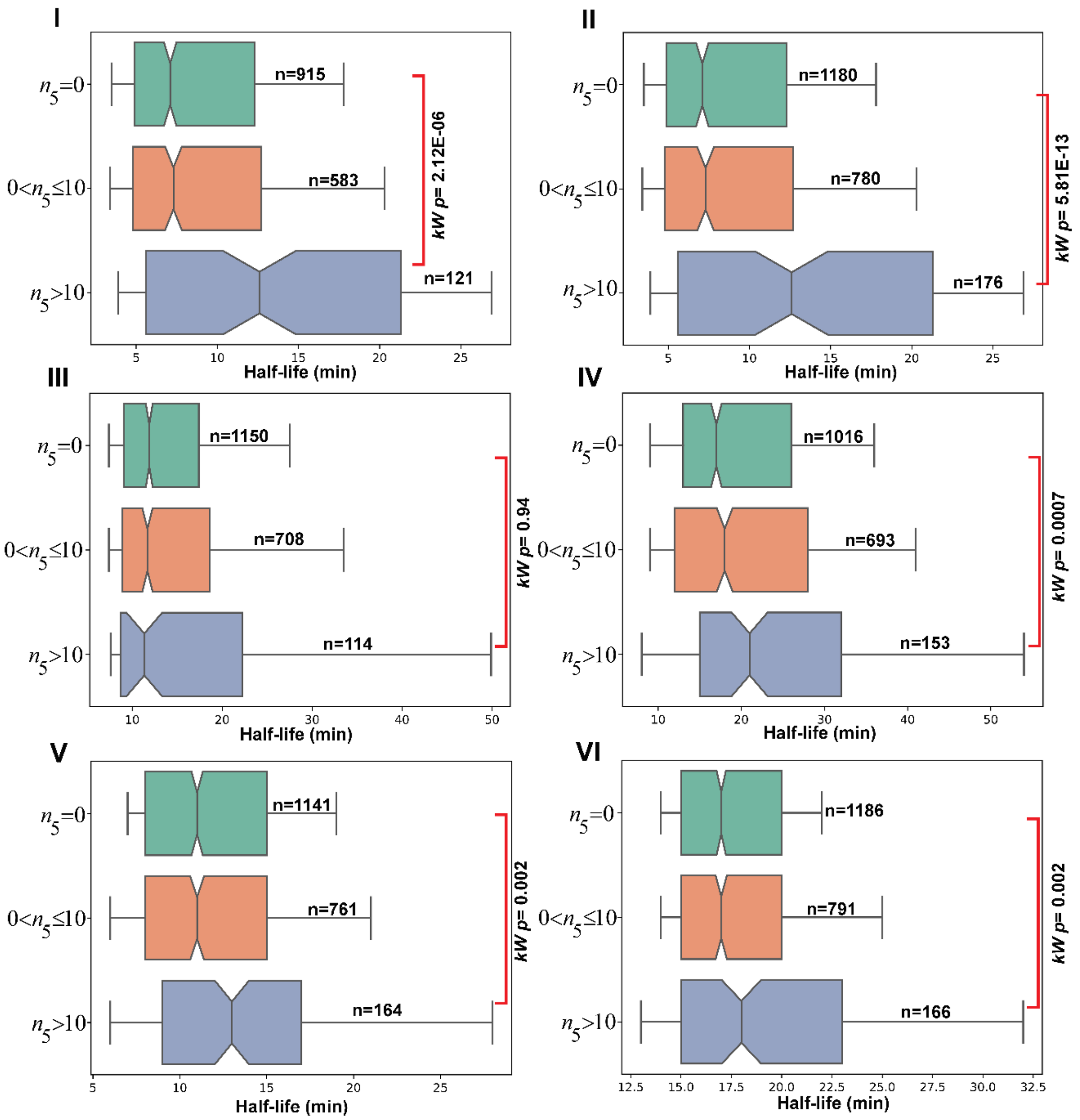


**Fig S4B.** Half-life distributions of *S. cerevisiae* mRNAs based on the number of unique proteins ($n_5$) binding to 5′ UTR for different datasets (I) Coller, (II) Cramer, (III) Gresham, (IV) Wang1, (V) Wang2, and (VI) Weis. These distributions are compared by a Kruskal-Wallis test, $p$-values are provided. In all these comparisons, same as Fig S4A, higher the number of bound proteins, longer is the half-life of the respective transcript.

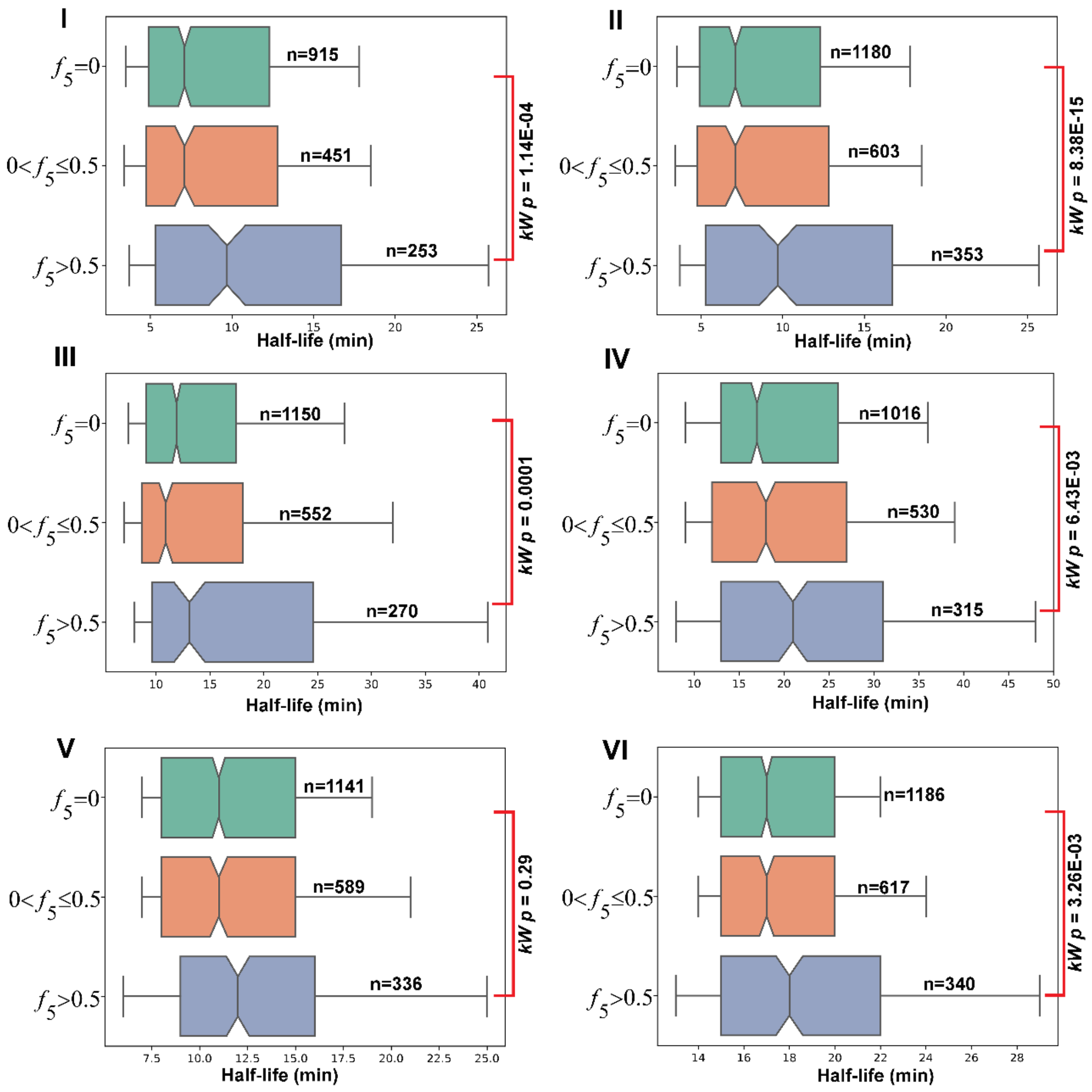


**Fig S4C.** Half-life distributions of *S. cerevisiae* mRNAs based on the fraction of 5′ UTR length covered by protein binding regions for different datasets (I) Coller, (II) Cramer, (III) Gresham, (IV) Wang1, (V) Wang2, and (VI) Weis. These distributions are compared by a Kruskal-Wallis test, *p*-values are provided. Increasing fraction of the 5′ UTR length covered by protein binding regions result in longer transcript half-life, presumably by hindering progressive 5′→3′ exonuclease digestion.

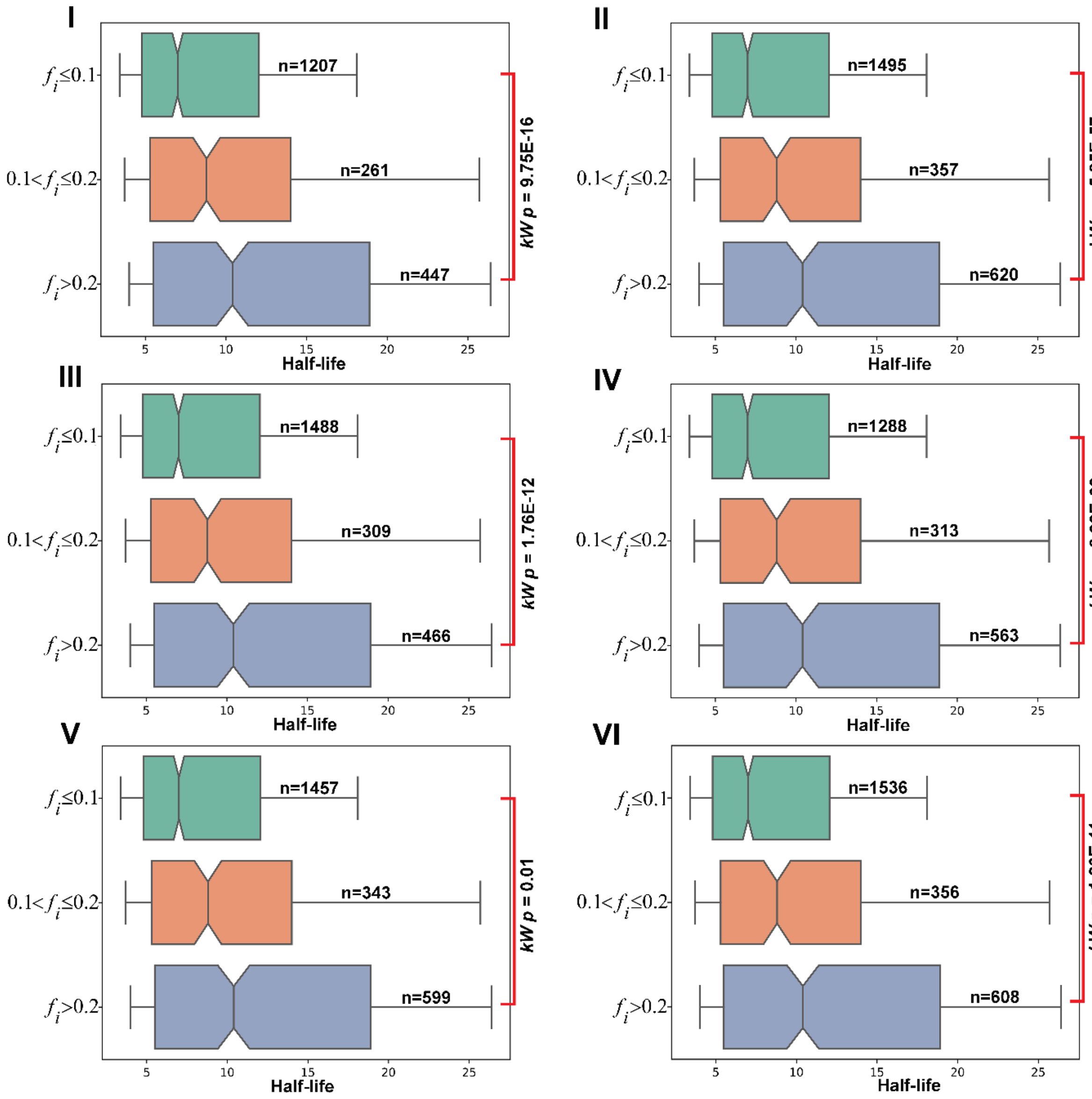


**Fig S4D.** Half-life distributions of *S. cerevisiae* mRNAs based on the fraction of internal unstructured region covered by protein binding regions for different datasets (I) Coller, (II) Cramer, (III) Gresham, (IV) Wang1, (V) Wang2, and (VI) Weis. These distributions are compared by a Kruskal-Wallis test, *p*-values are provided. Increased amount of internal unstructuredness within transcript showed decreased half-life. Burial of these internal unstructured segments by proteins would hinder easy engagement of the degradation machinery, and thus, would elevate the transcript half-life. For analysis of the internal unstructured region, we used a 12/6 window size (i.e., a window of 12 nt was sliding from 5′ to 3′ end, 6 nt slide at a time) and a window was assigned as a unstructured region if 60% nucleotides within the window had PARS < 0.

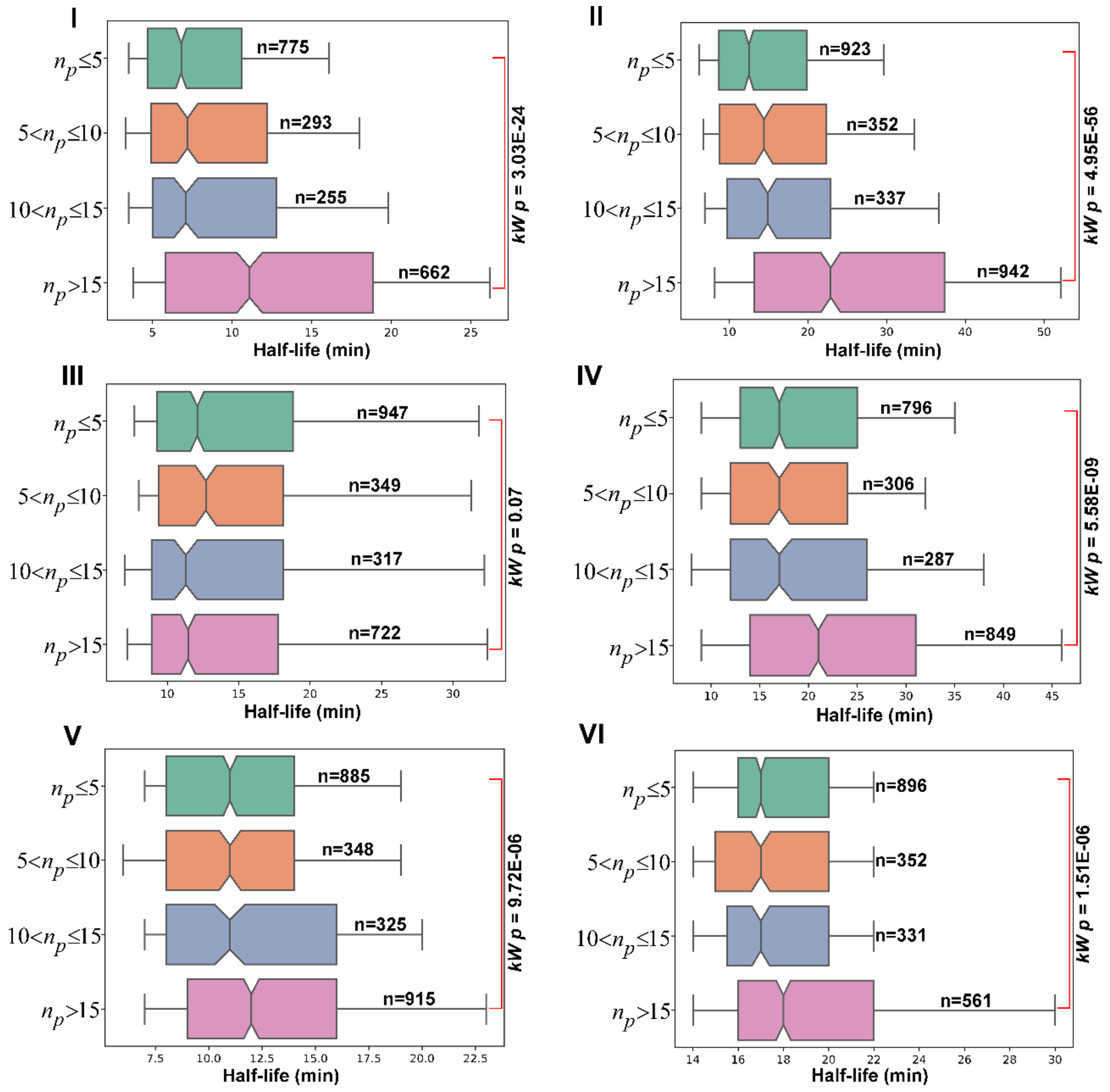


**Fig S4E.** Half-life distributions of *S. cerevisiae* mRNAs based on the total number of unique proteins binding to *S.cerevisiae* mRNA for different datasets (I) Coller, (II) Cramer, (III) Gresham, (IV) Wang1, (V) Wang2, and (VI) Weis. These distributions were compared by Kruskal-Wallis tests, *p*-values are provided. In these comparisons, increasing number of RBP binding results in longer transcript half-life.

### Supplementary Figure related to Figure 5 of main text

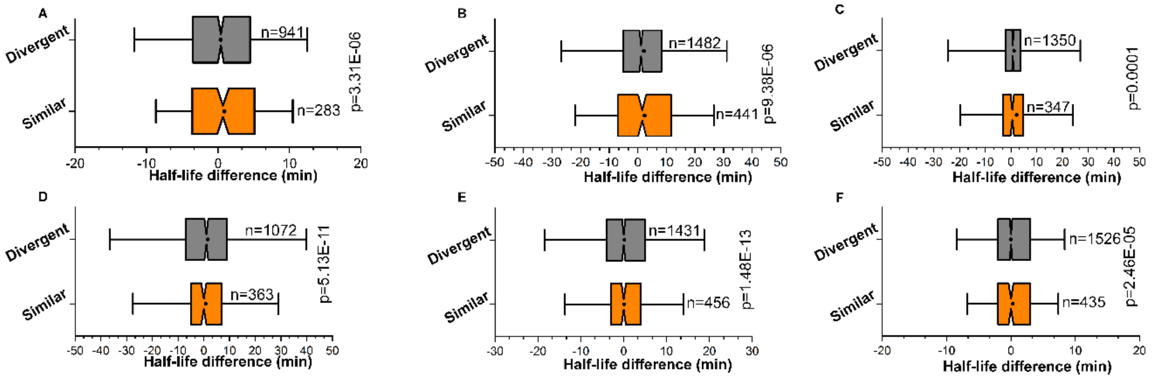


**Fig S5A.** Distribution of half-life differences in *S. cerevisiae* paralogous transcripts, grouped according to the difference in the 5′ and 3’ TUR lengths between the paralogous pair for different datasets (I) Coller, (II) Cramer, (III) Gresham, (IV) Wang1, (V) Wang2, and (VI) Weis. Similar: 5′ TURs of both transcripts are either ≤ 5 or > 5, as well as 3′ TURs are also ≤ 33 or > 33. Divergent: 5′ TUR of one paralog is ≤ 5, while the other is > 5, and 3′ TUR of one paralog is ≤ 33, while the other is > 33. These distributions are compared by a Mann-Whitney U test, *p*-values are provided.

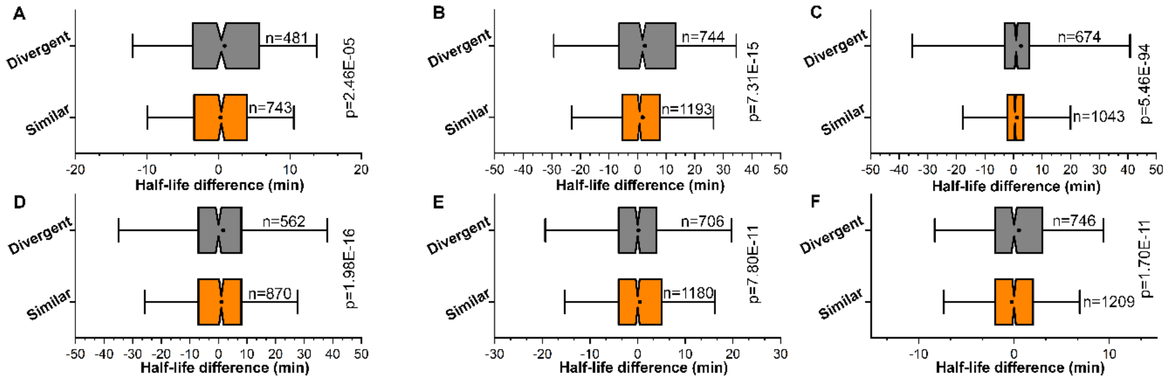


**Fig S5B.** Distribution of half-life differences in *S. cerevisiae* paralogous transcripts, grouped according to the difference in the IUS ($\xi$-value) between the paralogous pair for different datasets (I) Coller, (II) Cramer, (III) Gresham, (IV) Wang1, (V) Wang2, and (VI) Weis. Similar: $-0.05 \leq \Delta\xi \leq 0.05$. Divergent: $-0.05 > \Delta\xi$ or $\Delta\xi > 0.05$. These distributions are compared by a Mann-Whitney U test, *p*-values are provided.

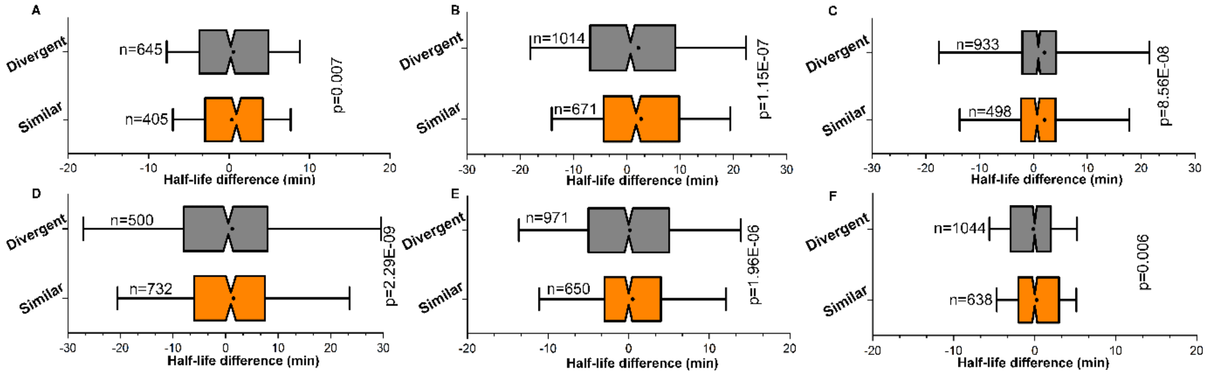


**Fig S5C.** Distribution of half-life differences in *S. cerevisiae* paralogous transcripts, grouped according to the difference in the RBP between paralogous pair for different datasets (I) Coller, (II) Cramer, (III) Gresham, (IV) Wang1, (V) Wang2, and (VI) Weis. Similar: those that bind to roughly the same number of proteins, and Divergent: those that bind to differential number of proteins. These distributions are compared by a Mann-Whitney U test, *p*-values are provided.
